# Structure of the Dengue Virus RNA Promoter

**DOI:** 10.1101/2022.04.15.488410

**Authors:** Yi-Ting Sun, Gabriele Varani

**Affiliations:** Department of Chemistry, University of Washington, Seattle, WA 98195-1700 (USA)

**Keywords:** NMR, Flaviviruses, Dengue virus, UTR, Stem loop A

## Abstract

Dengue virus, a single-stranded positive sense RNA virus, is the most prevalent mosquito-borne pathogen in the world. Like all RNA viruses, it uses conserved structural elements within its genome to control essential replicative steps. A 70 nucleotides stem-loop RNA structure (called SLA) found at the 5’-end of the genome of all flaviviruses, functions as the promoter for viral replication. This highly conserved structure interacts with the viral polymerase NS5 to initiate RNA synthesis. Here we report the NMR structure of a monomeric SLA from Dengue virus serotype 1, assembled to high-resolution from independently folded structural elements. The DENV1 SLA has an L-shape structure, where the top and side helices are coaxially-stacked and the bottom helix is roughly perpendicular to them. Because the sequence is highly conserved among different flavivirus genomes, it is likely that the three-dimensional fold and local structure of SLA are also conserved among flaviviruses and required for efficient replication. This work provides structural insight into the Dengue promoter and provides the foundation for the discovery of new antiviral drugs that target this essential replicative step.

## Introduction

Flaviviruses such as dengue (DENV), West Nile (WNV), yellow fever (YFV) and Zika virus (ZIKV), cause severe human diseases. Among them, Dengue fever is the most prevalent mosquito-borne viral disease in humans (Clyde et al., 2006; Gubler, 2006). It is estimated that up to 400 million people become infected with it each year (Murray et al., 2013). Any of the four dengue virus serotypes (DENV1 to DENV4) can produce clinical symptoms ranging from a flu-like syndrome to the severe and even fatal Dengue hemorrhagic fever (Clyde et al., 2006; Gubler, 2006). Despite the significant impact of Dengue infection on human health, effective vaccines are not yet available after > 70 years of efforts (Dyer, 2017), and small molecule treatment is only beginning to show promise (Lin et al., 2017; Raut et al., 2015; Saleem et al., 2019).

Dengue, like all flaviviruses, is a single-stranded positive sense RNA virus with a genome of about 11 kb, with a 5’-type I cap but without a polyadenylated tail (Colavita et al., 2020; Dang et al., 2020; Kamau et al., 2019; Pascalis et al., 2020; Selisko et al., 2014). The viral genome encodes a long polyprotein which is subsequently processed by both host and viral proteases to generate 10 mature viral proteins. Three structural proteins, Capsid (C), Envelop (E), and prM, assemble new viral particles, while seven non-structural proteins (NS1, NS2A, NS2B, NS3, NS4A, NS4B, and NS5) are responsible for viral RNA replication (Garcia-Blanco et al., 2016; Klema et al., 2015; Mazeaud et al., 2018; Neufeldt et al., 2018; Ng et al., 2017). The NS5 protein consists of an N-terminal methyltransferase (MTase) domain which is involved in capping, and a C-terminal RNA-dependent RNA polymerase (RdRp) domain which is responsible for RNA synthesis (Ferrer-Orta et al., 2006; Neufeldt et al., 2018; Yap et al., 2007; Zhao et al., 2015).

The coding sequence is flanked by 5’- and 3’-UTRs which contains highly conserved cis-acting RNA elements that are important for translation of viral proteins, RNA synthesis, encapsidation, and genome dimerization (Gebhard et al., 2011; Ng et al., 2017). In particular, the highly conserved stem-loop A (SLA) in the 5’-UTR, close to the 5’-end of the genome, functions as a “promoter’ for the NS5 protein to initiate minus-strand synthesis (Choi, 2021; Filomatori et al., 2006; Gebhard et al., 2011). The interaction between SLA and NS5 is critical for RNA synthesis and was confirmed to be direct by biochemical studies; removal of SLA and mutation of certain nucleotides drastically decreases RNA replication (Filomatori et al., 2006; Lodeiro et al., 2009). Consistent with its essential functional role, the sequence and secondary structure of the SLA are highly conserved (Figure 1).

**Figure 1.**
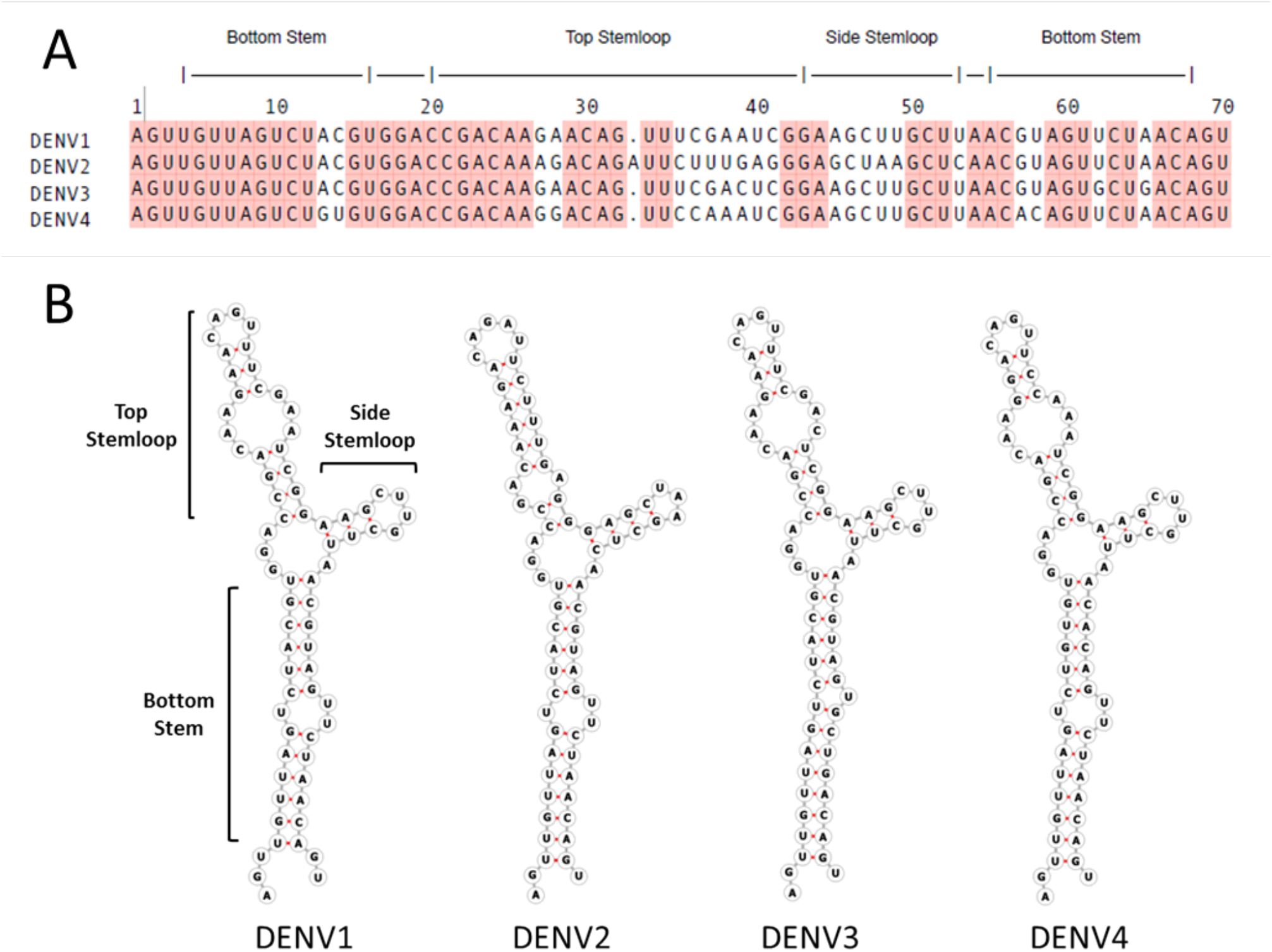
The sequence and secondary structure of the SLA promoter element that recruits the NS5 protein to initiate minus-strand RNA synthesis is very highly conserved among the four Dengue virus serotypes (DENV1-4). (A) Alignment of the sequences of the first 70 nucleotides of DENV1, DENV2, DENV3, and DENV4. The regions corresponding to the predicted secondary structure elements (bottom helix, top and side stem-loops) are indicated at the top. Conserved nucleotides are shaded. (B) Predicted RNA secondary structure of the SLA for the four DENV serotypes, as confirmed by SHAPE (Dethoff et al., 2018).

The current model for DENV RNA synthesis involves cyclization of the genome, mediated by long-range RNA-RNA interactions through inverted complementary sequences near the opposite ends of the RNA, which position the 5’- and 3’-UTRs in close proximity (Alvarez, Lodeiro, et al., 2005; Filomatori et al., 2011; Filomatori et al., 2006; Villordo et al., 2010). This interaction allows the SLA-bound NS5 to transfer to the 3’-stem loop (3’-SL) at the end of the 3’-UTR and initiate minus-strand synthesis (Hodge et al., 2016; Mazeaud et al., 2018). Removal of the SLA or 3’-SL leads to a significant reduction of viral replication (Alvarez, De Lella Ezcurra, et al., 2005; Filomatori et al., 2006; Yu et al., 2008), suggesting that a small molecule targeting these RNAs could have anti-viral activity.

In all flaviviruses, the SLA forms a Y-shaped secondary structure, consisting of a bottom helix with a conserved U-rich bulge, a short side stem-loop and an apical stem-loop containing an internal loop (Figure 1) (Dethoff et al., 2018; Gritsun & Gould, 2007). A crystal structure of DENV2 SLA was determined using a chimeric RNA, wherein the DENV2 SLA was inserted into the anticodon loop of a human tRNA (Lee et al., 2021). It revealed a large L-shape dimer, where the side stem-loop of each monomer is base paired with another molecule in the crystal. It is unclear whether dimerization is an artifact of crystallization, or a relevant functional state, perhaps transient, of the virus.

Here we report the NMR structure of the monomeric DENV1 SLA. Because of the severe peak overlap caused by the large size of the RNA, we use a divide-and-conquer approach to assemble a high-resolution structure from three independently folded structural elements corresponding to two of the three stem-loops, and the three-way junction (Figure 2), which were then assembled to generate the full SLA RNA through NMR analysis of the complete element (Barnwal et al., 2016). The SLA has an L-shape structure, consistent with the crystal structure, where the top and side stem-loops are coaxially-stacked and the bottom helix is roughly perpendicular to it. We only observe formation of a dimeric conformation at NMR concentrations, above 0.5 mM; suggesting that if a dimer does indeed form in the cell, it is not because of its thermodynamic stability. Our structure will facilitate understanding of the mechanism of dengue virus replication and provides the foundation for the discovery of new antiviral drugs, which is already under way in the group.

**Figure 2.**
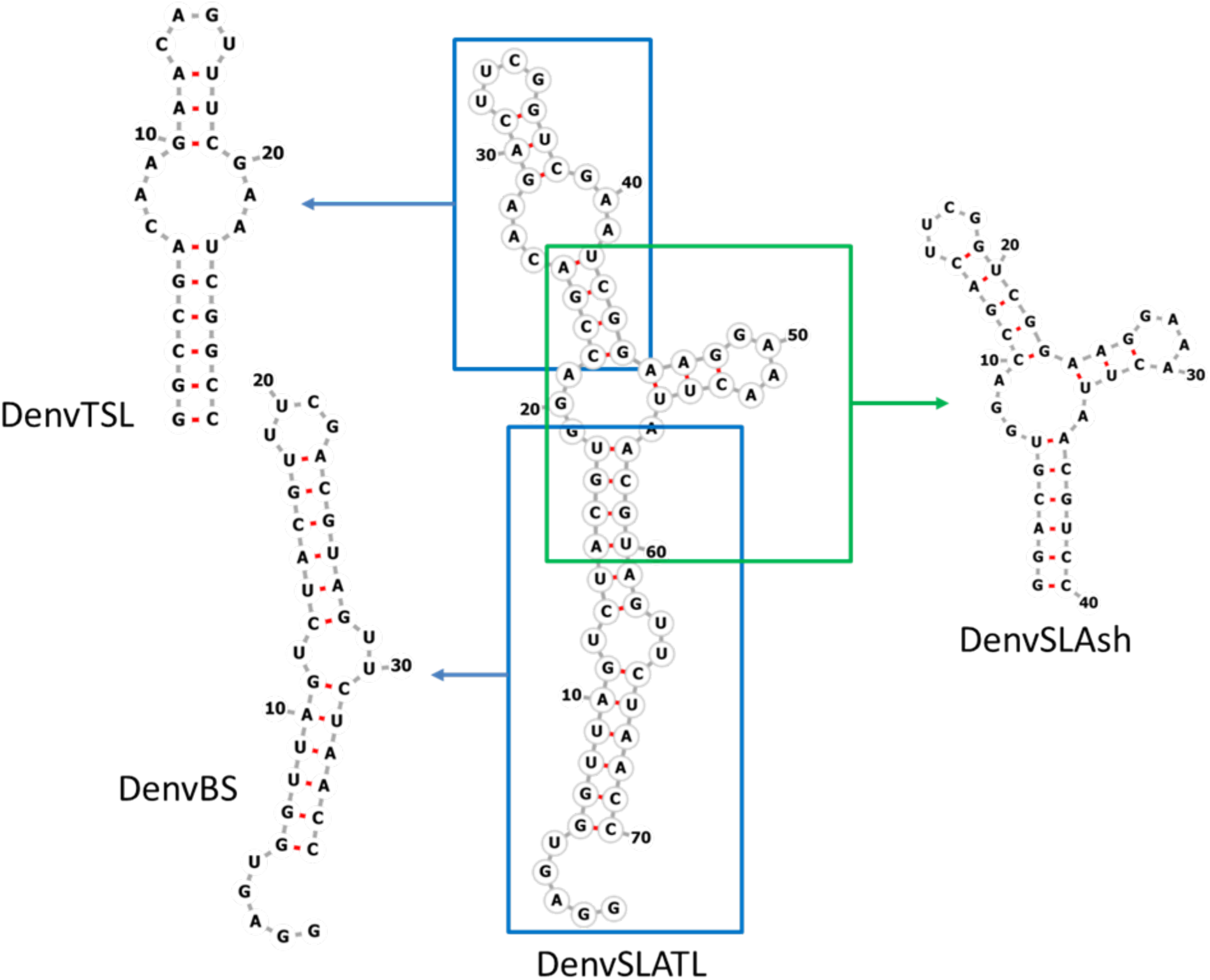
The sequences and secondary structures of each RNA segment studied in this work. Three smaller constructs, corresponding to independently folded secondary structure domains, were prepared to facilitate structure determination: DenvBS represents the bottom helix; DenvTSL the top stem-loop and DenvSLAsh the three-way junction. Secondary structures for each segment were predicted using the UNAFold web server (http://www.unafold.org) (Markham & Zuker, 2008; Zuker, 2003) and verified by NMR.

## Results

### Construct design

The spectra of the complete SLA RNA are characterized by extensive spectral overlap and broad lines because of its relatively large size, 70 nts, causing difficulties in obtaining unambiguous peak assignment and, especially, collecting a large number of constraints for structure determination (Supplementary Figure S1). Thus, we adopted the strategy of reconstructing the complete structure by analyzing smaller independently folded structural elements (divide-and-conquer) (Barnwal et al., 2016). This can be done because RNA structure is modular, and its folding hierarchical, and secondary structural elements generally fold independently outside of the context of the complete RNA (Tinoco & Bustamante, 1999). This is very different from proteins, where secondary and tertiary structure folding generally cannot be separated. Nevertheless, the validity of this approach has to be experimentally verified in each case, which we do as described below.

We divided the DENV1 SLA into three structural segments that overlap to generate the complete promoter: the bottom helix (DenvBS), the top stem-loop (DenvTSL) and the three-way junction (DenvSLAsh) (Figure 2 and Supplementary Figure S2). Additional G-C base pairs were added at the end of the sequences to improve *in vitro* transcription and stabilize the local secondary structure, if needed. Tetraloops were added to some RNA sequences as well to cap the structure and stabilize it. Thus, a UUCG tetraloop was added to the top of DenvBS, DenvSLAsh, and the apical loop of DenvTSL. For the complete SLA, we incorporated UUCG and GAAA tetraloop into the top and side stem-loop of DENV1 SLA as well, respectively, to generate a stabilized SLA which we named DenvSLATL (Figure 2, Supplementary Figure S1 and S2), and to avoid dimer formation through the side stem-loop, as described next.

### The structure of the individual domains recapitulates what is observed in the full SLA

The divide-and-conquer approach is only warranted if the structure of individual domains faithfully recapitulates what is observed in the complete RNA. For each of the smaller structural segments, imino proton peaks of base-paired residues predicted from the secondary structure were observed in H2O NOESYs, except for fast exchanging imino protons for unpaired nucleotides and the base-pairs at the end of helical stretches (Supplementary Figure S3). Assignments of these imino protons allow us to rapidly establish and verify the predicted secondary structures for each domain.

High quality NMR spectra could be collected for a tetraloop-stabilized SLA model that avoid dimerization through the side stem-loop (see below, DenvSLATL) and peak assignment was facilitated by the assignments of each segment. Reassuringly, we find that each NH in the smaller segments has similar chemical environment to its corresponding NH in the complete RNA, thus presenting similar chemical shifts and verifying that the structure observed in the complete SLA is retained in the individual fragments. Namely, overlaying the NOESY NMR spectra of the segments on the DenvSLATL spectra revealed high similarities in chemical shifts and NOE patterns (Figure 3), which confirms the secondary structure of the fragments coincide with what is seen, for the corresponding domain, in the complete RNA.

**Figure 3.**
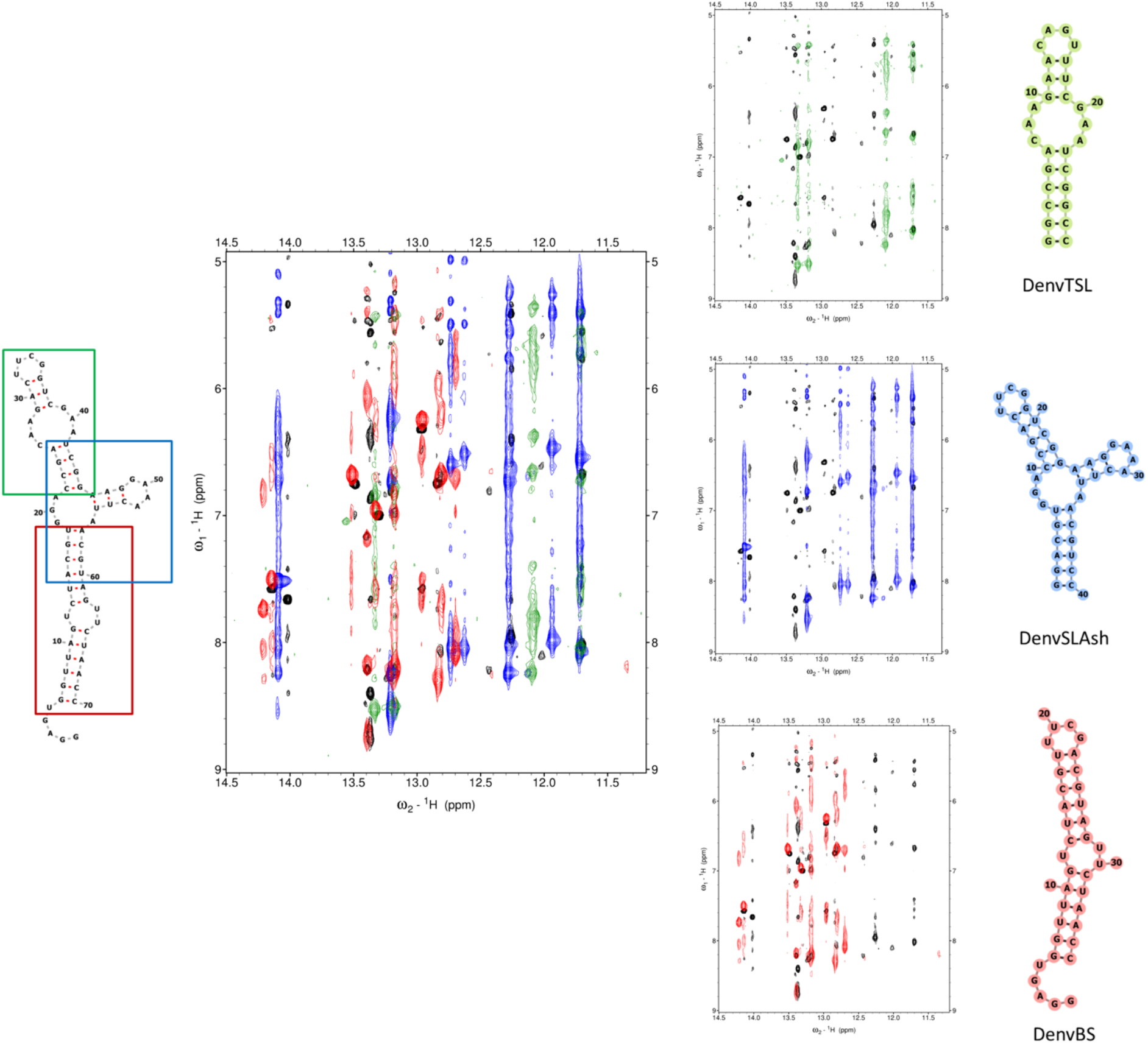
Overlay of the imino region of NOESY spectra for DenvSLATL (black) and the three smaller constructs which were prepared to facilitate structure determination. Black color corresponds to DenvSLATL, red to DenvBS, the bottom helix; green corresponds to DenvTSL, the top stem-loop, and blue corresponds to DenvSLAsh, the three-way junction. 1H NOESY spectra were recorded in 10 mM potassium phosphate buffer (pH 6.5, 90% H2O/ 10% D2O) at 25°C. Despite small differences, high similarity of chemical shifts and NOE patterns were observed, allowing the transfer of peak assignments from each segment to the SLA model oligonucleotide (Supplementary Figure S3).

NOESY spectra in D2O were then collected to assign non-exchangeable protons. Extensive overlaps in the sugar proton region were relieved by deuteration of H3’, H4’, H5’, H5’’ and H5 protons (Supplementary Figure S4-S6), even if this approach prevented assignments of most of those protons. ^15^N- and ^13^C-labeled samples were prepared for DenvTSL to record ^1^H-^15^N HSQC and ^1^H-^13^C HSQC to distinguish ambiguous peaks in NOESY spectra, as well. Through deuteration, we were able to observe characteristic sequential NOEs in helices and assign non-exchangeable protons for each of the separate Denv structural segments (Supplementary Figure S4-S6).

The existence of a single dominant and monomeric conformation was confirmed in each case from the number of Ura and Cyt H5-H6 peaks in TOCSY spectra, which was in all cases consistent with the sequence (Supplementary Figure S7). Formation of base-paired double helices were confirmed by cross-strand NOEs involving imino resonances, as well as sequential NOEs involving both exchangeable and non-exchangeable protons (Supplementary Figure S4-S6). Thus, DenvBS and DenvTSL were confirmed to form stem loop structures as expected; the monomeric/dimeric state of the three-way junction is discussed below.

### NMR Structures of the individual structural elements of DENV1 SLA RNA

Once assignments were completed, NOE distance constraints were systematically tabulated for structure determination, and the constrain list was refined by multiple rounds of structure calculations. For each of the three structural segments, structure calculations were performed independently using distance and torsion angle restraints derived from NMR experiments, as summarized in the Supplementary Tables S1-S3. Once the segments were completed, restraints from corresponding nucleotides in each segment were also added to the restraint table for structure calculation of the complete monomeric SLA (see below).

RNA structures were calculated with Xplor-NIH using torsion angle dynamics and simulated annealing starting from randomized coordinates against the restraint table (Schwieters et al., 2003). A total of 1,000 independent calculations were executed and the lowest energy structure was further refined by an extended simulated annealing calculation against the experimental restraints. 400 structures were eventually generated, and the 10 lowest scoring structures were selected for presentation for each RNA molecule. The resulting structures converge well, with no NOE restraint violation greater than 0.5 Å. Representative structures for DenvSLA segments were presented with PyMol (Figure 4) and the overall heavy atom RMSDs are listed in Supplementary Tables S1-S3 as well.

**Figure 4.**
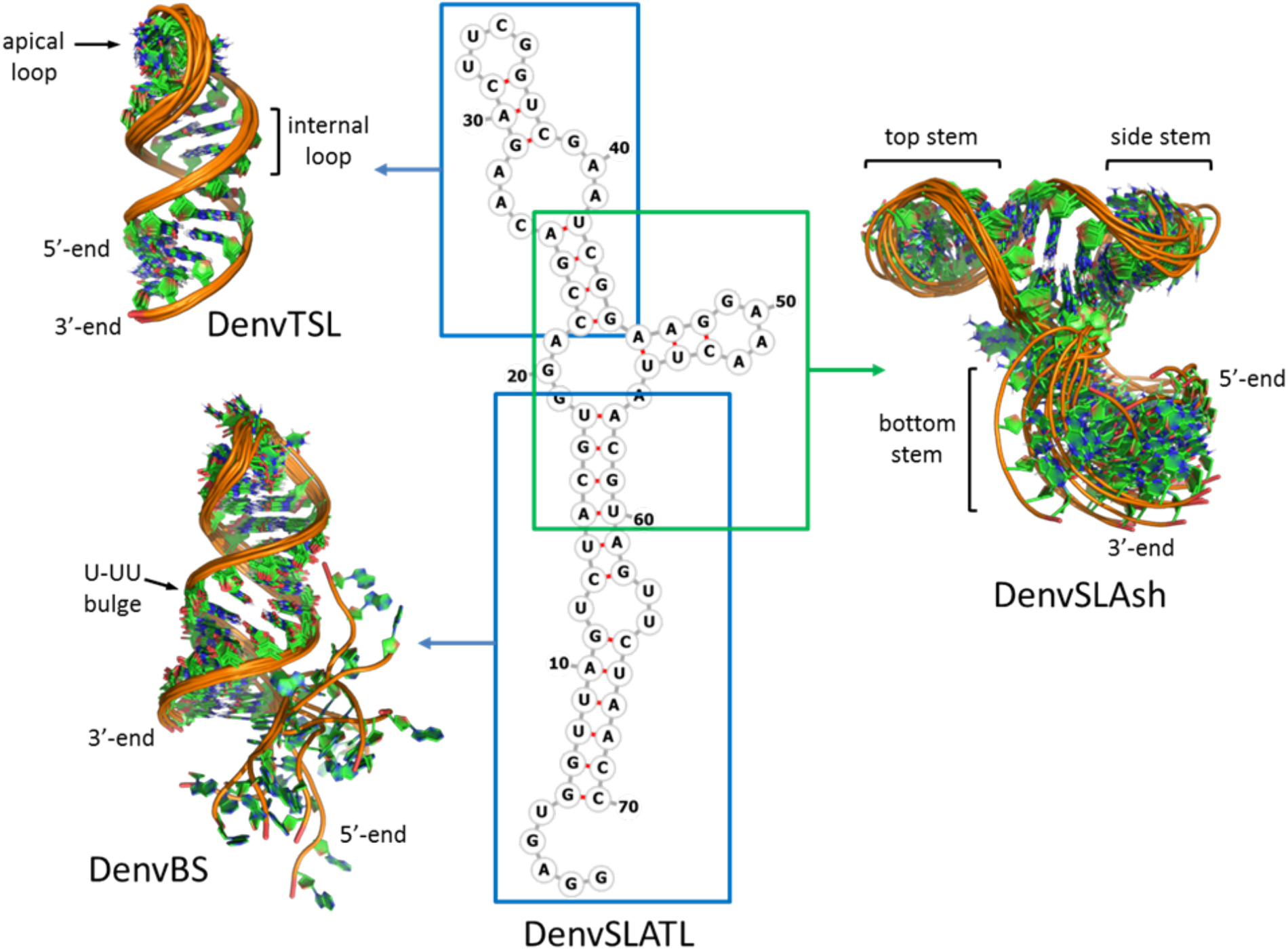
Superposition of the 10 NMR structures of lowest energy for each independently folded RNA segment studied in this work. DenvBS represents the bottom helix and forms a well-defined double helix with an unstructured 5’-tail (G1-U5) and a U-rich bulge; DenvTSL represents the top stem-loop and forms a stable helix interrupted by an internal loop, capped by a partially flexible loop; DenvSLAsh represents the three-way junction and forms a L-shape structure, where the top and side stem helices stack coaxially, and the bottom helix is roughly perpendicular to the coaxial stack; this last image was generated by superposing only the two stacked helices to emphasize the coaxial stack and partial flexibility at the 3- way junction.

As predicted, DenvBS and DenvTSL form double helical stems interrupted by internal loops. In both cases, no direct evidence of non-canonical base-pair formation in the internal loops was found, but sequential aromatic NOE correlations (H6/H8-H6/H8) in those regions were observed (Supplementary Figure S4 and S5), together with other sugar-base NOEs, implying the retention of helical stacking throughout the internal loop. Many of the unpaired nucleotides are therefore stacked inside the helices, and potentially base paired, but we lack the information to impose such constraints, perhaps because any non-canonical base pairs are only transiently formed.

The resulting structures provide insight into the local structure in these regions. In the U-rich internal loop of DenvBS, Ura12 and Ura30 are coplanar and Ura29 stacks between Gua28 and Ura30 (Figure 5A); nucleotides in the internal loop of DenvTSL (Cyt7 to Ade9 and Gua20 to Ade22) also stack within the helical stem, retaining coaxial stacking (Figure 5B). Although the terminal loop of DenvTSL varies somewhat within the 10 calculated structures, the topology is clearly established. The Gua15 base points toward the major groove, while Ade14 and Ura16 points outward, in the direction of the solvent, and could conceivably provide direct interactions with the RdRP or other accessory proteins (Figure 5C). The significance of each region in Dengue SLA had been validated by biological studies and is discussed further in the Discussion Section below.

**Figure 5.**
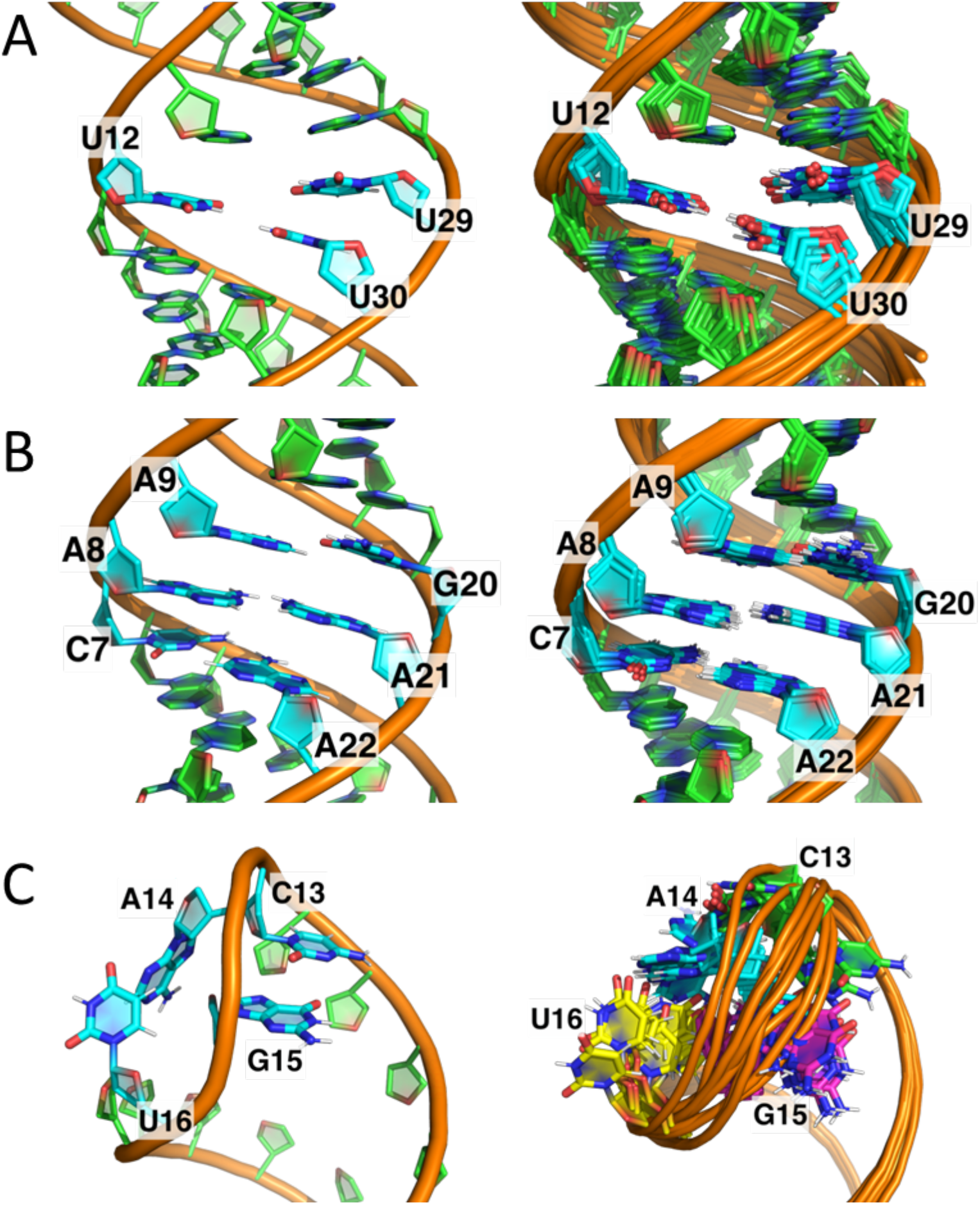
(A) Close-up view of the U-UU bulge in DenvBS, drawn from the lowest energy structure (left) and superposition of 10 representative NMR structures (right). Ura12 and Ura30 are coplanar, potentially forming an unstable base pair (no NH peak is visible in 2D spectra), while Ura29 stacks between Gua28 and Ura30, retaining continuous stacking. (B) A close-up view of the internal loop in DenvTS, taken from the lowest energy structure (left) and the superposition of 10 representative NMR structures (right). Nucleotides Cyt7-Ade9 and Gua20-Ade22 across the internal loop stack within the helix. Although no evidence of non-canonical base-pairs formation in this region was observed in NMR spectra, uninterrupted sequential NOEs were observed (Supplementary Figure S5). (C) A close-up view of the terminal loop in DenvTSL. from the lowest energy structure (left) and the superposition of 10 representative NMR structures (right). In this last image, nucleotides are colored differently for clarity (C13 in green, A14 in cyan, G15 in magenta, and U16 in yellow). In all 10 structures, Gua15 points toward the major groove while Ade14 and Ura16 point outward towards the solvent, while Cyt13 is poorly defined.

### Structure of the 3-way junction

The key structural element in SLA is the 3-way junction which organizes the complete element; because a dimer, is observed in the DENV2 structure, it remains unclear what the three-dimensional organization of the full SLA would be. In order to investigate the three-way junction in DENV1 SLA with the necessary resolution, we prepared an RNA containing the sequence of the three-way junction and side stem-loop, but with shortened top and bottom helical stems to reduce spectra overlap. Furthermore, two RNAs were synthesized, one containing the wild-type side stem loop (SLAshCUUG) and the other containing a GAAA-stabilized side stem loop (DenvSLAsh) (Figure 2 and Supplementary Figure S1). This was done because we observed extra base pairs in the NMR spectra of SLAshCUUG. The signal intensity for those extra base pairs is concentration dependent, which indicates a monomer-dimer equilibrium at the 0.5-1.5 mM concentration of our NMR experiments (Supplementary Figure S8A). By substituting CUUG with GAAA tetraloop, dimerization was eliminated (Supplementary Figure S8B).

We presume that with a less stabilizing loop sequence, the side stem loop in SLAshCUUG would open and form dimers through RNA-RNA interactions at the mM concentrations of NMR experiments (Supplementary Figure S8C), as observed in the crystal. Considering the low copy number of viral RNAs in cell, it is very unlikely that the dimer is the thermodynamically favored SLA structure and therefore we collected NMR spectra of the monomeric DenvSLAsh for structural analysis. However, we cannot exclude the possibility that dimer formation occurs at intermediate steps of viral replication; this is further discussed below.

Base-pair imino NOEs and characteristic sequential NOE patterns for helical stem nucleotides were observed in NOESY spectra for the three double helical stems (Supplementary Figure S3 and S6), validating the secondary structure prediction. For nucleotides Gua7, Gua8 and Ade9, only sequential NOEs were observed, but no other inter-stranded or inter-nucleotide NOEs were identified. NOE interactions corresponding to Gua23H8-Ade24H8, Ura33H6-Ade34H8 and Ade34-Cyt10H6 were observed in NOESY spectra as well (Figure 6A), consistent with continuous stacking between the top and side stem-loops, and suggestive of coaxial stacking of the two helices.

**Figure 6.**
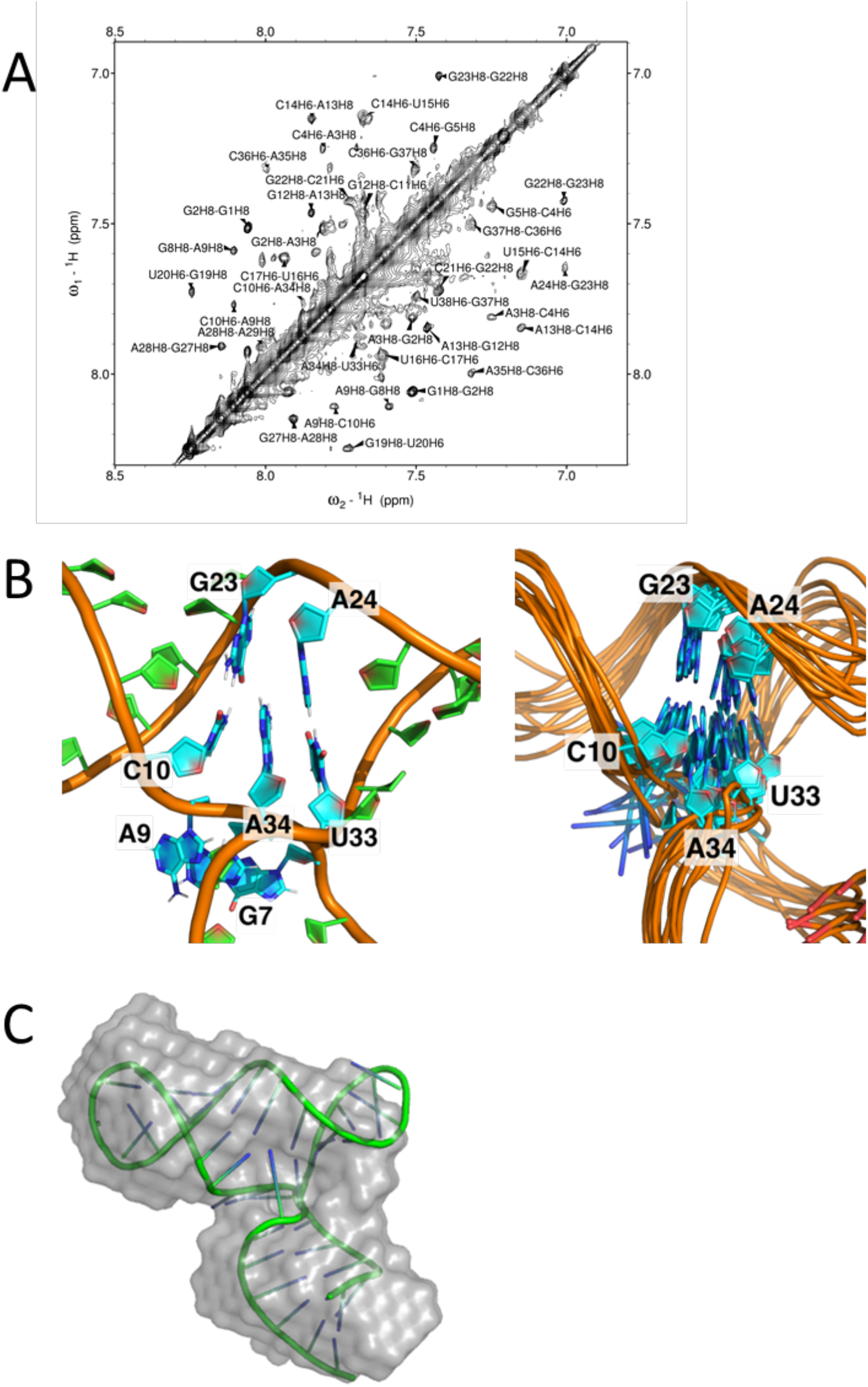
(A) NOESY spectrum of DenvSLAsh showing non-exchangeable H6/H8-H6/H8 correlations. This spectrum was recorded at 25°C in 10 mM potassium phosphate buffer (pH 6.5, 90% H_2_O/ 10% D_2_O). The following NOE interactions: G23H8-A24H8, U33H6-A34H8 and A34-C10H6 in the three-way junction establish coaxial stacking of the top and side stem-loops. (B) A close-up view of the three-way junction in DenvSLAsh from the lowest energy structure (left) and superposition of 10 representative NMR structure (right). Ade34 stacks within the coaxial helix, between Cyt10 at the end of top stem and Ura33 at the end of the side stem-loop. (C) The lowest energy structure of DenvSLAsh in shown in cartoon representation superposed on the SAXS model; the SAXS results were not used for NMR refinement, and therefore provide independent validation of the structure.

The calculated structure of DenvSLAsh demonstrates formation of an L-shape three-way junction, where the top and side stem loops in stack coaxially, and the bottom helix is roughly perpendicular to the coaxial stack (Figure 4). Ade34 in the three-way junction stacks within the coaxial helix, between Cyt10 in the top stem and Ura33 in the side stem. Gua7, Gua8 and Ade9 are instead flexible and single stranded, giving the three-way junction some flexibility, and thus allowing the angle of the bottom helix to wiggle relative to the top coaxial stack (Figure 6B).

The topology of the three-way junction structure was independently validated by SAXS analysis (Figure 6C and Supplementary Figure S9). Both SLAshCUUG and DenvSLAsh exhibit L-shape SAXS envelops which agree very well with the L-shape NMR structure and provide independent validation. They also demonstrate that the structure is fully monomeric below at least 0.1 mM concentration.

### Structure of the Denv1 promoter

By examining each of the independently folded secondary structure elements separately, we were able to establish the local structures for the two stem-loops and 3-way junction and side stem-loop that form Denv1 SLA. Spectral and structural information collected from these spectra were used to assist peak assignments and structure determination for the complete SLA structure.

A single conformation of DenvSLATL was confirmed by the number of Ura and Cyt H5- H6 peaks in TOCSY spectra (Supplementary Figure S10). Overlay of the NOESY and TOCSY spectra of DenvSLATL with those of its structural segments revealed very similar chemical shifts and NOE patterns (Figure 3 and Supplementary Figure S10) which allowed us to safely transfer the much larger number and more confidently assigned restraints obtained from the individual segments. Structure calculations were then performed for DenvSLATL using distance and torsion angle restraints derived from experimental NMR data of DenvSLATL and each of the structural segments, as was done for each of the elements in the structure. The 10 lowest structures converge with no NOE restraint violation greater than 0.5 Å and high structural precision, and SAXS was then used to validate the structure. Importantly, the topology of the structure of DenvSLATL was independently validated by SAXS analysis, and the SAXS envelope agree very well with the L-shape NMR structure (Figure 7). The structural statistics are summarized in Supplementary Table S4.

**Figure 7.**
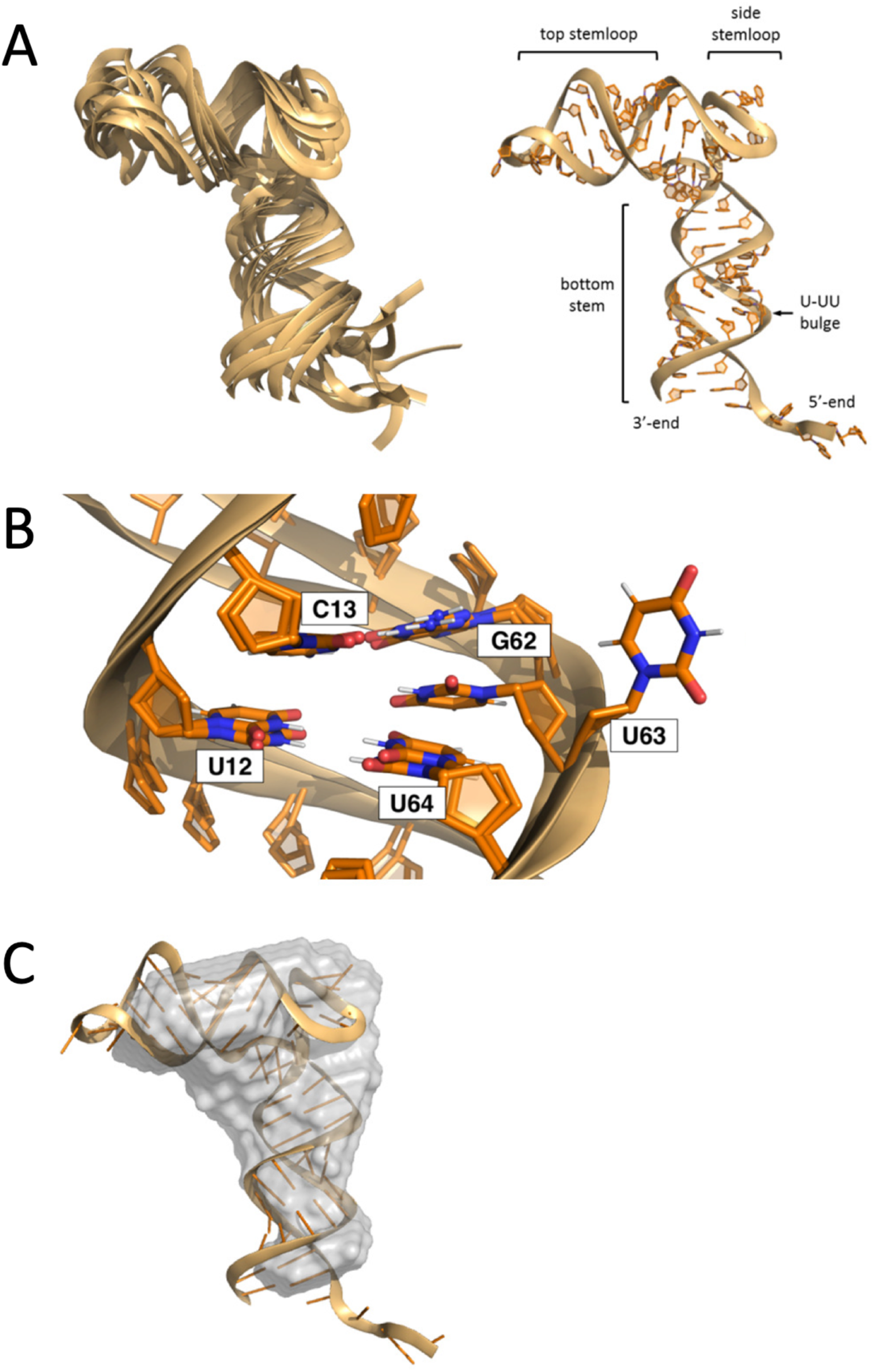
(A) Superposition of the 10 NMR structures of DenvSLATL (left), and the structure with the lowest calculated energy (right). (B) A close-up view of the U-rich bulge of DenvSLATL. Two conformations of U-UU bulge were observed in the calculated NMR structures. (C) The lowest energy structure of DenvSLATL in cartoon representation superposed on the SAXS envelope; the SAXS results were not used for refinement, and therefore provide independent validation of the NMR structure.

The calculated structure of DenvSLATL reveals an L-shape RNA, where the top and side stem loop are coaxial and the bottom stem roughly perpendicular to it (Figure 7A), as per the smaller three-way junction. The arrangement of nucleotides in the three-way junction is as observed in DenvSLAsh: Ade56 stacks between the base-pairs of Cyt22-Gua45 and Ade46-Ura66, which are at the end of the top and side stem-loop, respectively, while nucleotides in the longer internal junction loop (Gua19 to Ade21) have some flexibility and thus the orientation of the bottom stem is less well defined. The U-rich bulge in the bottom stem is important for SLA function. In our NMR structures, Ura12 and Ura64 are co-planar and stack within the helix. However, two conformations of Ura63 were observed in the calculated structures: one with the base within the helix and another with the base pointing outward (Figure 7B). Eight out of the lowest 10 structures have Ura63 stacked within the helix and two structures have Ura63 pointing outward, suggesting conformational flexibility.

## Discussion

The Denv SLA structure functions as an RNA “promoter’ for the viral RNA-dependent RNA polymerase enzyme NS5 and is essential viral replication. NS5 is recruited to the viral RNA through this element, which has high conservation in both sequence and secondary structure across flaviviruses (Choi, 2021; Filomatori et al., 2006; Gebhard et al., 2011). The presence of a U-rich bulge in the bottom helix is essential for SLA function, and the high conservation of the 3-way junction supports a functional role for the overall 3D shape of the RNA, which is determined by the topology of the junction. We observe a rigid conformation for the three-way junction, with co-axial stacking of the top and side stem-loops, to generate a well-defined L-shape structure, with the bottom helix emanating at a nearly 90 angle, but with some flexibility because of the single stranded nucleotides linking the bottom and top helices (Figure 7A). It is possible that the conformational flexibility we observe plays a role in its function, since RNA-binding proteins often exploit induced fit (Leulliot & Varani, 2001), but this remains to be investigated.

In our NMR structure, Ura63 in the U-rich bulge occupies 2 conformations; the base can point inward or outward, but the 2 conformations do not affect the rest of the RNA structure (Figure 7B); in the recently reported crystal structure of a dimeric Denv2 SLA, the same Uracil points outward (Lee et al., 2021). The fact that DENV3 has a single U bulge in the bottom stem and mutational studies of the U-UU bulge imply that Ura63 is critical for SLA promoter function, but the two remaining nucleotides are not. Deletion or mutations of the U-rich bulge largely impair viral RNA replication *in vivo*, but the effect on the binding affinity to RdRp and its *in vitro* activity is insignificant (Filomatori et al., 2011; Lodeiro et al., 2009). This suggests that U-rich bulge interacts with another protein which is important for viral replication in infected cells.

The apical loop is important for SLA function and highly conserved; the CAG(X)U sequence is found in all four Dengue serotypes (Figure 1), and mutations impair both viral replication *in vivo* and RdRp activity *in vitro* (Filomatori et al., 2011; Filomatori et al., 2006; Lodeiro et al., 2009). Interestingly, mutations in the terminal loop do not significantly affect RdRp binding, suggesting the terminal loop might play an important role in post-binding steps to promote polymerase activity (Filomatori et al., 2011). In the NMR structure, we observe that Gua32 points towards the major groove and faces the other two loop nucleotides Ade31 and Ura33, which point outward (Figure 5C). The orientation of the loop nucleotides could be important for RdRp activity. The top stem, however, is less conserved among dengue serotypes. I*n vivo* biological assays have shown that the presence of an internal loop in the top stem has no significant effect on SLA function either (Lodeiro et al., 2009), yet shortening the helix significantly decreases viral replication and RdRp binding (Filomatori et al., 2011; Lodeiro et al., 2009). This implies that, although the helical stem is unlikely to provide direct interactions with the NS5 protein, it is important to space the terminal loop relative to the three-way junction and the U-rich bulge.

The three-way junction is likely to provide a structural framework to orient the SLA and different domains of NS5, giving it an L-shaped conformation confirmed by the SAXS analysis. The top and side stem-loop are coaxially stacked, as expected, since no single-stranded nucleotide is found at the junction between them (Lescoute & Westhof, 2006). The stacking of Ade56 in the linker between side and bottom stem constrains the direction of the bottom stem to be roughly perpendicular to the coaxially stacked helix.

The recently reported crystal structure of Denv2 SLA also presents a L-shape structure, but this is generated by dimerization in the crystal created by the unpaired side loop which forms RNA-RNA kissing loop interactions (Lee et al., 2021). In the crystal structure, the side loop was open and engaged in loop-loop interactions. We observe two conformations for a construct containing the wild-type loop sequence, consistent with monomer-dimer equilibrium. Replacement of the wild-type loop with GAAA stabilizes the monomer and allowed us to establish the structure of a monomeric SLA RNA. It remains to be seen whether SLA dimerization is of functional relevance. Because we observe increased dimer peak intensity at the NMR concentrations, 0.5-1.5 mM, and no dimer below 0.5 mM, we presume that under cellular conditions, the SLA RNA will be entirely monomeric. However, this conclusion does not preclude the possibility that the dimeric structure is transiently present as a result of protein binding or dimerization of the genome, and in fact it could be an elegant way for the virus to regulate its promoter activity.

We expect the 3D structure of the three-way junction to be conserved in flaviviruses (Figure 1 and Supplementary Figure S12), because no nucleotides are predicted to be present in the junction between top and side stem in all flavivirus sequences, and the linker between side and bottom stem is always short, while the longest linker occurs invariably between bottom and top helices. It follows that the shape of the SLA and the orientation of the top and bottom stems are likely important for SLA to be recognized by the viral polymerase.

In summary, we have established the 3D structure of the thermodynamically favored, monomeric form of the SLA promoter from Dengue virus serotype 1; which is very likely to be representative of all other serotypes, and indeed it is very likely all flaviviruses will share the same global structural arrangement. In addition to providing a framework for interpreting biochemical data and NS5 activity, this structure also lays the groundwork to identify small molecule inhibitors that target the highly conserved 3-way junction, which are being actively pursued in our group.

## Materials and Methods

### RNA Preparation

All RNA molecules were synthesized by in vitro transcription with *in house* purified T7 RNA polymerase using synthetic DNA oligonucleotide templates (purchased from Integrated DNA Technologies) and standard methods (Milligan et al., 1987). Partially deuterated RNA molecules (deuteration of H5, H3’, H4’, H5’ and H5’’ protons) were synthesized using selective deuterated NTPs (from Cambridge Isotopes). ^15^N- and ^13^C-labeled samples were synthesized, if needed, using isotope-labeled NTPs (from Cambridge Isotopes). The sequence and secondary structures of all the RNAs, as verified by NMR, are shown in Supplementary Figure S2.

The RNA oligonucleotides were purified by denaturing polyacrylamide gel electrophoresis (PAGE), electroeluted and concentrated by ethanol precipitation (Gubser & Varani, 1996). After extensive dialysis into 10 mM potassium phosphate buffer (pH 6.5), the RNAs were annealed by heating briefly to 90 °C followed by snap cooling in an ice bath. Final RNA concentrations used for NMR studies were 0.6∼1.2 mM. For experiments studying non-exchangeable protons, samples were lyophilized to dryness and dissolved into D2O. Samples used to study exchangeable protons were dissolved in H2O:D2O (9:1).

### NMR Spectroscopy

All NMR spectra were collected at 25°C on Avance III 600 MHz, AVANCE III 700MHz or AVANCE III 800MHz spectrometer equipped with cryogenic probes (600 and 800). The 1D ^1^H spectra were recorded using the excitation sculpting pulse sequence. 2D total correlation spectroscopy (TOCSY) spectra were recorded with mixing times of 80 ms. The exchangeable and non-exchangeable 2D NOESY spectra were recorded with various mixing times (100, 200 and 300 ms) to assist spectral assignments and quantitative evaluation of internuclear distances by comparison with peak intensities for pair of protons with fixed distances. Spectra for selective deuterated samples were collected in the same manner. 2D ^1^H-^15^N and ^1^H-^13^C HSQC spectra were recorded on isotope labeled samples, if needed to confirm assignments. All NMR data were processed with TOPSPIN (Bruker) and analyzed in NMRFAM-SPARKY (Lee et al., 2015). Assignments of RNA spectra were guided by predicted RNA chemical shift values and based on well-established double-helical sequential NOE patterns (Varani et al., 1996; Varani & Tinoco, 1991).

### Experimental Restraints and Structure Determination

Interproton distance restraints were derived from NOE cross-peaks in 2D 1H NOESY spectra and sorted into strong (2.5 ± 0.7 Å), medium (3.5 ± 1.2 Å) and weak (4.5 ± 1.5 Å) bins based on peak intensities, relative to fixed distances (e.g. H5-H6 = 2.5 Å, H3’-H6/H8 = 3.5 Å). Base-pair planarity and hydrogen-bonding restraints were used for unambiguously established base pairs as identified by 2D NOESY involving NH protons. Hydrogen bond, planarity and dihedral restraints were included for base-paired nucleotides that were surrounded by base pairs confirming to A-form helical structures, as established from the pattern of NOE cross-peaks (Varani et al., 1996; Varani & Tinoco, 1991).

We have often found within the group that flexible loops capping stem-loops lead to loss of spectral quality, most likely due to non-specific aggregation multimerization at NMR concentrations (Barnwal et al., 2016; Varani et al., 1991); this was the case for this RNA as well, with the added complication of dimerization through kissing loop interactions which created multiple conformations. Thus, UUCG and GAAA tetraloops were used to replace the dynamic apical loop or added to the end of the bottom helix to improve spectra quality (Banas et al., 2010; Jucker & Pardi, 1995; Varani et al., 1991).

Experimental constraints for structure calculation of the complete DenvSLATL were compiled by dividing the RNA into three segments: DenvBS, DenvTSL and DenvSLAsh (Figure 2), corresponding to the bottom helix, apical stem-loop and three-way junction and side stem-loop. Overlay of 2D ^1^H-^1^H NOESY spectra from the segments, with a spectrum of the complete Dengue 1 SLA model (called DenvSLATL), showed strong similarities in the chemical shifts, which allowed the transfer of segments assignments to DenvSLATL for assignments for structure calculations (Figure 3). Restraints derived from DenvBS spanned nucleotides G1-U18 and A57-A70; those obtained from DenvTSL spanned nucleotides C22- G45; DenvSLAsh contributed information around the three-way junction, namely A15-A25 and U42-U60; obviously, there is overlap which allowed us to further verify that the divide-and-conquer approach was warranted. The NMR experimental constraints are summarized in Supplementary Tables S1-S4.

RNA structures were calculated with Xplor-NIH using torsion angle dynamics and simulated annealing from a single extended RNA starting template (Schwieters et al., 2003). Compiled experimental restraints were used in a simulated annealing procedure, initially undergoing high-temperature (2,500K to 298K) torsional angle dynamics, where incremental decreases in temperature were generated by progressively introducing Van der Waals terms and increasing force constants for angles, dihedral angles, NOEs, and the Van der Waals repulsive term. After the final cooling step, the RNA underwent two sequential refinement steps, first in torsional angle space then in Cartesian space. A total of 1,000 independent calculations were executed for each RNA. and the lowest energy structure was further refined by an extended simulated annealing calculation against the constraints listed above to generate 400 structures. The 10 lowest scoring structures were then examined in PyMol, and structure quality analysis was conducted using MolProbity. Representative structures of DenvBS, DenvTSL, DenvSLAsh and DenvSLATL were deposited in Protein Data Bank with accession codes: 7K4L, 7UME, 7UMD and 7UMC, respectively.

### Small-angle X-ray Scattering (SAXS)

RNA samples for SAXS were prepared similarly to samples made for NMR spectroscopy, but at different concentrations. Samples with varied concentrations (1-5 mg/mL) were prepared in 20 mM Tris, 100 mM NaCl and 0.1 mM EDTA (pH 6.5). SAXS experiments were recorded on an *in-house* state-of-art SAXS instrument (BioSAXS-2000) at Argonne National Laboratory. The data were processed using RAW and particle distance distribution function P(r) plots were calculated using GNOM (Svergun, 1992) and used for low resolution *ab initio* shape reconstruction with DAMMIN (Svergun, 1999). A total of 20 models were generated with DAMMIN using the ATSAS online server (https://www.embl-hamburg.de/biosaxs/) (Manalastas-Cantos et al., 2021). The representative model was selected with a suite of software tools (DAMSEL, SAMSUP, DAMAVER and DAMFILT) for comparison and fitting to the NMR structure (Volkov & Svergun, 2003).

DAMSEL compares the models, finds the most probable model and identifies outliers; DAMSUP aligns all models with the most probable model; DAMAVER averages these aligned models and computes a probability map; and DAMFILT filters the average model at a given default cut-off volume, which is the expected volume of the generated PDB file. DAMFILT removes loosely defined and lower occupancy atoms and generates a most probable compact model.

## Acknowledgments

We wish to thank all members of the Varani group for discussion and support; Dr. Greg Olsen for help with final preparation of the manuscript. The study was supported by NIH grant 1 R35 GM126942. This project has also been funded in part with Federal funds from the National Institute of Allergy and Infectious Diseases, National Institutes of Health, Department of Health and Human Services, under Contract No. HHSN272201700059C.

We acknowledge the use of SAXS Core facility of Center for Cancer Research, National Cancer Institute (NCI) of National Institutes of Health (NIH). The SAXS core resource has been funded in whole or in part with federal funds from NCI under contract 75N91019D00024 and the Intramural Research Program of the NIH, National Cancer Institute, Center for Cancer Research. The content of this publication does not necessarily reflect the views or policies of the Department of Health and Human Services, nor does mention of trade names, commercial products, or organizations imply endorsement by the U.S. Government. The SAXS data were collected at beamline 12-ID-B of Advanced Photon Source (APS) of Argonne National Laboratory (ANL). We thank Dr. Yu-Xing Wang (NCI) and Dr. Lixin Fan (FNLCR/Leidos) and Dr. Xiaobing Zuo (ANL) for their support. Use of the APS was supported by the U. S. Department of Energy, Office of Science, Office of Basic Energy Sciences, under Contract No. DE-AC02-06CH11357.

## Author Contributions

Yi-Ting Sun: Conceptualization, methodology, investigation, writing, reviewing and editing; graphics. Gabriele Varani: Experimental design, data analysis, writing, reviewing and editing.

