## Supplemental File for "Structure of the Dengue Virus RNA Promoter"

*Supplementary Table S1.* NMR experimental restraints and structural statistics for the 10 lowest energy structures of DenvBS (bottom helix)

| | DenvBS |
|---|---|
| **NMR Peak Assignment (excluding the unstructured 5'-tail, nucleotides 1-5)** | |
| All H1′, H2′, H5, H6, H8, AdeH2, 87.5% of H3', 59.4% of H4' and 18.8% of H5'/H5" were assigned. Extensive overlap in the sugar proton region was relieved by deuteration of H3', H4', H5', H5 "and H5 protons to assign H1' and H2' unambiguously, but this prevented assignments of many of those sugar protons. 63.2% of imino protons and 85.7% of Cyt amino protons were assigned. | |
| **NMR Experimental Restraints** | |
| Total number of restraints | 539 |
| Total NOE Restraints | 299 |
| Intra-residue | 162 |
| Inter-residue | 137 |
| Sequential li-jl = 1 | 116 |
| Non-sequential li-jl > 1 | 21 |
| Hydrogen bond restraints | 10 |
| Dihedral Angle Restraints | 216 |
| Planarity Restraints | 24 |
| **Structure Analysis** | |
| NOE violations >0.5 Å | 0 |
| Torsion angle violations >5° | 0 |
| **Deviation from Idealized Geometry** | |
| Bond lengths (Å) | 0 |
| Bond angles (°) | 0 |
| **Heavy Atom RMSD to the Mean Structure (Å)** | |
| All RNA heavy atoms (6-36, excluding the unstructured 5'-tail) | 1.02 |
| All RNA backbone (6-36, excluding the unstructured 5'-tail) | 1.00 |
| RNA upper stem heavy atoms (13-28) | 0.72 |
| RNA upper stem backbone (13-28) | 0.60 |
| RNA lower stem heavy atoms (6-12, 29-36) | 0.50 |
| RNA lower stem backbone (6-12, 29-36) | 0.53 |

*Supplementary Table S2.* NMR experimental restraints and structural statistics for the 10 lowest energy structures of DenvTSL (apical stem-loop)

| | DenvTSL |
|---|---|
| **NMR Peak Assignment** | |
| All H1′, H2′, H5, H6, H8, 25% of AdeH2[(a)], 92.9% of H3', 67.9% of H4' and 37.5% of H5'/H5'' were assigned. Extensive overlap in the sugar proton region was relieved by deuteration of H3', H4', H5', H5 ''and H5 protons to assign H1' and H2' unambiguously, but this prevented assignment of those deuterated sugar protons. 58.3% of imino protons and 75% of Cyt amino protons were assigned. | |
| **NMR Experimental Restraints** | |
| Total number of restraints | 475 |
| Total NOE Restraints | 251 |
| Intra-residue | 135 |
| Inter-residue | 116 |
| Sequential Ii-jI = 1 | 92 |
| Non-sequential Ii-jI > 1 | 24 |
| Hydrogen bond restraints | 9 |
| Dihedral Angle Restraints | 206 |
| Planarity Restraints | 18 |
| **Structure Analysis** | |
| NOE violations >0.5 Å | 0 |
| Torsion angle violations >5° | 0 |
| **Deviation from Idealized Geometry** | |
| Bond lengths (Å) | 0 |
| Bond angles (°) | 0 |
| **Heavy Atom RMSD to the Mean Structure (Å)** | |
| All RNA heavy atoms (1-28) | 0.85 |
| All RNA backbone (1-28) | 0.75 |
| RNA upper stem heavy atoms (7-22) | 1.03 |
| RNA upper stem backbone (7-22) | 0.84 |

(a) H2 of Adenines located in single-stranded loops were not assigned.

*Supplementary Table S3.* NMR experimental restraints and structural statistics for the 10 lowest energy structures of DenvSLAsh (three-way junction)

| | |
|---|---|
| DenvSLAsh | |
| **NMR Peak Assignment** | |
| 90% of H1′, 95% of H2′, 94.1% of H5, 97.5% of H6/H8, 47.5% of H3', 27.5% of H4', 16.3% of H5'/H5'' and 40% of AdeH2[(a)] were assigned. Extensive overlap in the sugar proton region was relieved by deuteration of H3', H4', H5', H5 ''and H5 protons to assign H1' and H2' unambiguously, but this prevented assignments of those deuterated sugar protons. 65% of imino protons and 80% of Cyt amino protons were assigned. | |
| **NMR Experimental Restraints** | |
| Total number of restraints | 840 |
| Total NOE Restraints | 562 |
| Intra-residue | 262 |
| Inter-residue | 300 |
| Sequential Ii-jI = 1 | 187 |
| Non-sequential Ii-jI > 1 | 113 |
| Hydrogen bond restraints | 41 |
| Dihedral Angle Restraints | 250 |
| Planarity Restraints | 28 |
| **Structure Analysis** | |
| NOE violations >0.5 Å | 0 |
| Torsion angle violations >5° | 0 |
| **Deviation from Idealized Geometry** | |
| Bond lengths (Å) | 0 |
| Bond angles (°) | 0 |
| **Heavy Atom RMSD to the Mean Structure (Å)** | |
| All RNA heavy atoms (1-40) | 2.75 |
| All RNA backbone (1-40) | 2.67 |
| RNA top and side stem heavy atoms (10-33) | 2.07 |
| RNA top and side stem backbone (10-33) | 1.88 |
| RNA bottom stem heavy atoms (1-9, 34-40) | 1.76 |
| RNA bottom stem backbone (1-9, 34-40) | 1.11 |

(a) H2 of Adenines located in single-stranded loop were not assigned.

*Supplementary Table S4.* NMR experimental restraints and structural statistics for the 10 lowest energy structures of DenvSLATL (full-length monomeric SLA)

| | DenvSLATL |
|---|---|
| **NMR Experimental Restraints** | |
| Total number of restraints | 1277 |
| Total NOE Restraints | 843 |
| Intra-residue | 402 |
| Inter-residue | 441 |
| Sequential \|i-j\| = 1 | 303 |
| Non-sequential \|i-j\| > 1 | 138 |
| Hydrogen bond restraints | 68 |
| Dihedral Angle Restraints | 384 |
| Planarity Restraints | 50 |
| **Structure Analysis** | |
| NOE violations >0.5 Å | 0 |
| Torsion angle violations >5° | 0 |
| **Deviation from Idealized Geometry** | |
| Bond lengths (Å) | 0 |
| Bond angles (°) | 0 |
| **Heavy Atom RMSD to the Mean Structure (Å)** | |
| All RNA heavy atoms (6-70) | 3.98[a] |
| All RNA backbone (6-70) | 4.08[a] |
| Top stemloop heavy atoms (22-45, corresponding to DenvTSL) | 1.33 |
| Top stemloop backbone (22-45, corresponding to DenvTSL) | 1.26 |
| Bottom stem heavy atoms (6-18, 57-70, corresponding to DenvBS) | 1.02 |
| Bottom stem backbone (6-18, 57-70, corresponding to DenvBS) | 1.00 |
| Three-way junction heavy atoms (15-25, 42-60, corresponding to DenvSLAsh) | 2.68 |
| Three-way junction backbone (15-25, 42-60, corresponding to DenvSLAsh) | 2.68 |

(a) The RMSD is calculated without the unstructured single-stranded 5'-tail (residues 1-5).

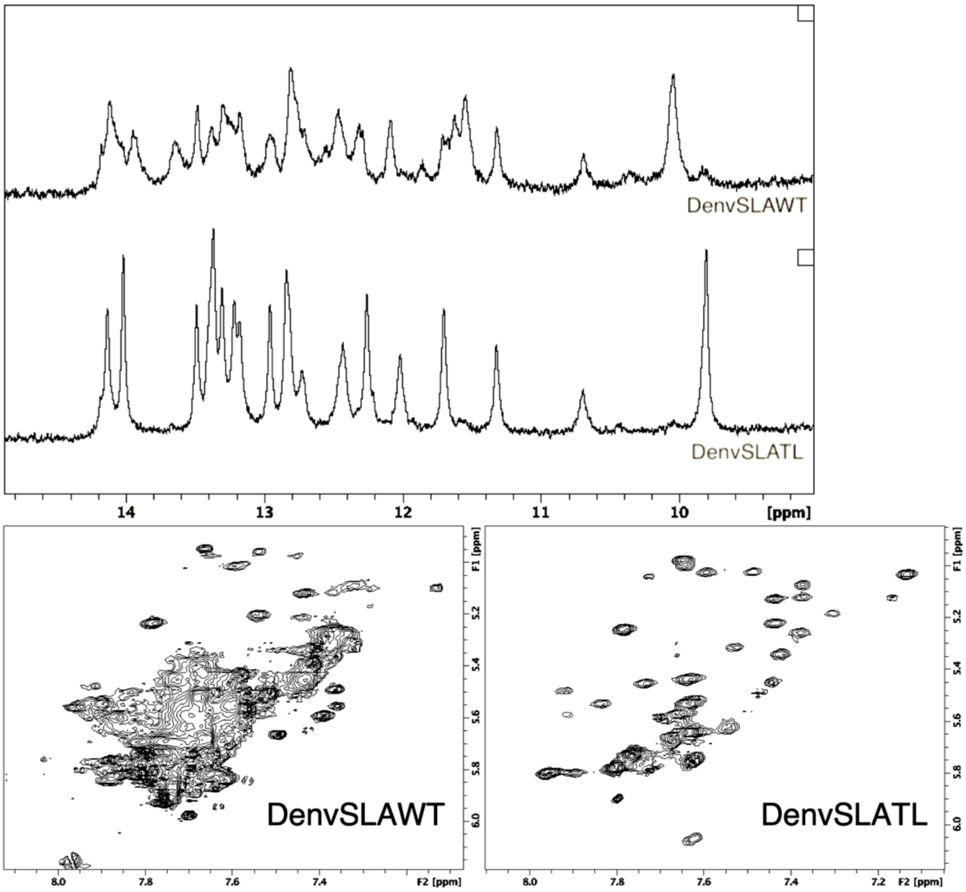


*Supplementary Figure S1.* (A) Comparison of the imino region of the 1D $^1$H spectra of wild-type DENV1 SLA (called DenvSLA-WT) and a tetraloop-stabilized SLA promoter (called DenvSL-ATL). The spectra are very similar, but the quality is much higher for the tetraloop-containing RNA, in part as a result of partial dimerization through the side stem-loop at the mM concentrations of NMR experiments, which would further increase the linewidth and the number of peaks in the spectrum. (B) Comparison of the H5-H6 region of the 2D TOCSY spectra of wild-type (DenvSLAWT) and tetraloop-stabilized SLA (DenvSLATL). While many of the peaks are in similar locations, the spectra for DenvSLAWT have much broader linewidth resulting in severe peak overlaps, which would make peak assignments for DenvSLAWT extremely difficult. The addition of tetraloops improves linewidth and allows identification of discrete peaks, as we have observed with multiple RNAs in the past (Barnwal et al., 2016; Sharma & Varani, 2020; Walker et al., 2020). This strategy is analogous to what is often done in crystallography, where crystallization modules (e.g. protein binding sites) are introduced to facilitate crystal packing and increase resolution.

| Name | Length | Sequence |
|---|---|---|
| DenvSLAWT | 70 | GGAGU UGUUA GUCUA CGUGG ACCGA CAAGA ACAGU UUCGA AUCGG AAGCU UGCUU AACGU AGUUC UAACA |
| DenvSLATL | 70 | GGAGU GGUUA GUCUA CGUGG ACCGA CAAGA CUUCG GUCGA AUCGG AAGGA AACUU AACGU AGUUC UAACC |
| DenvBS | 36 | GGAGU GGUUA GUCUA CGUUU CGACG UAGUU CUAAC C |
| DenvTSL | 28 | GGCCG ACAAG AACAG UUUCG AAUCG GCC |
| DenvSLAsh | 40 | GGACG UGGAC CGACU UCGGU CGGAA GGAAA CUUAA CGUCC |
| SLAshCUUG | 40 | GGACG UGGAC CGACU UCGGU CGGAA GCUUG CUUAA CGUCC |

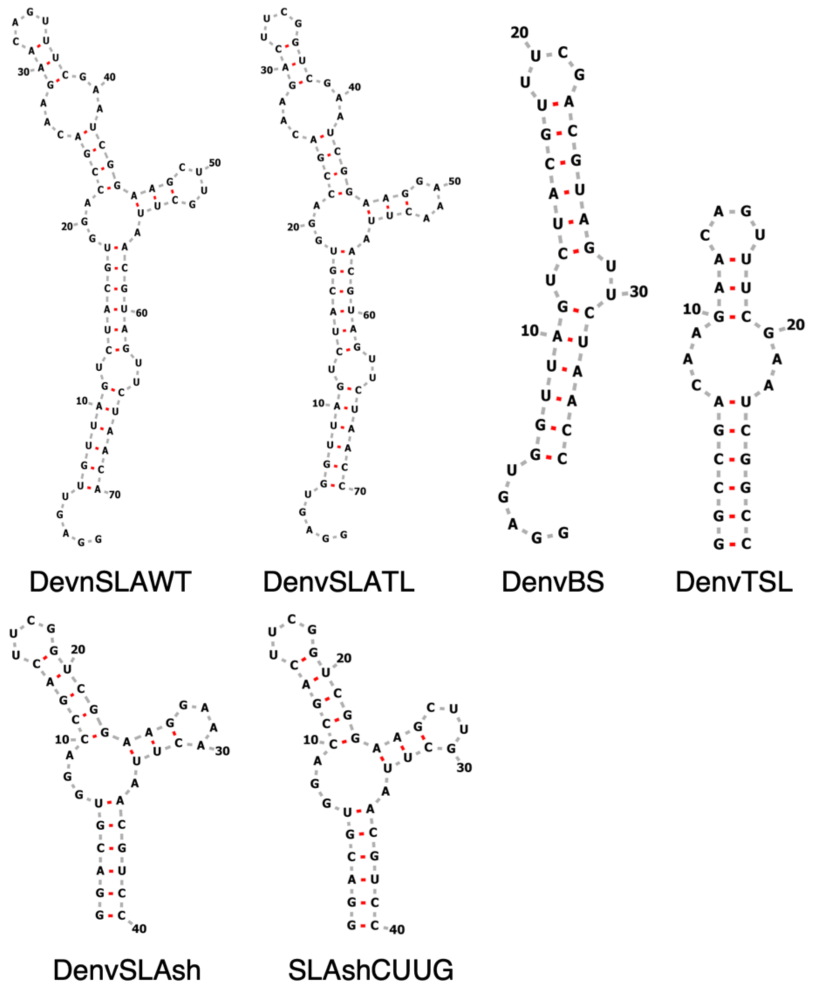


*Supplementary Figure S2.* Sequences and secondary structures of all RNAs studied in this work; all secondary structures were verified by NMR assignments of NH resonances.

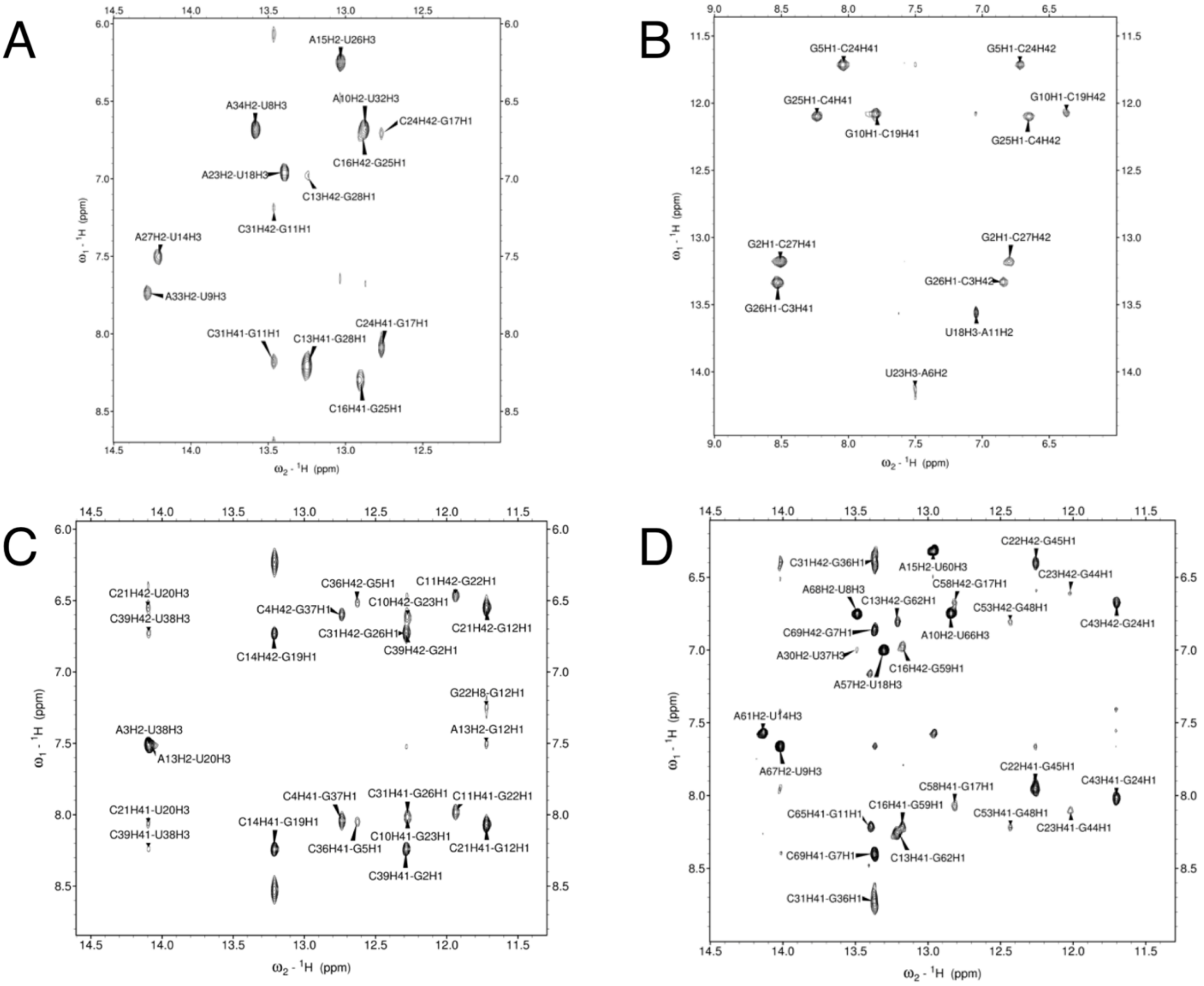


*Supplementary Figure S3.* Imino to amino/H2 regions of the NOESY spectra of (A) DenvBS, (B) DenvTSL, (C) DenvSLAsh and (D) DenvSLATL. All imino proton peaks of base-pairs predicted from the secondary structure of DenvBS, DenvTSL and DenvSLAsh were observed, except for fast exchanging imino protons for unpaired nucleotides and the base-pairs at the end of helices, validating the predicted secondary structure. Assignments for DenvSLATL were safely transferred from each corresponding segments. Spectra were recorded in 10 mM potassium phosphate buffer (pH 6.5, 90% H2O/ 10% D2O).

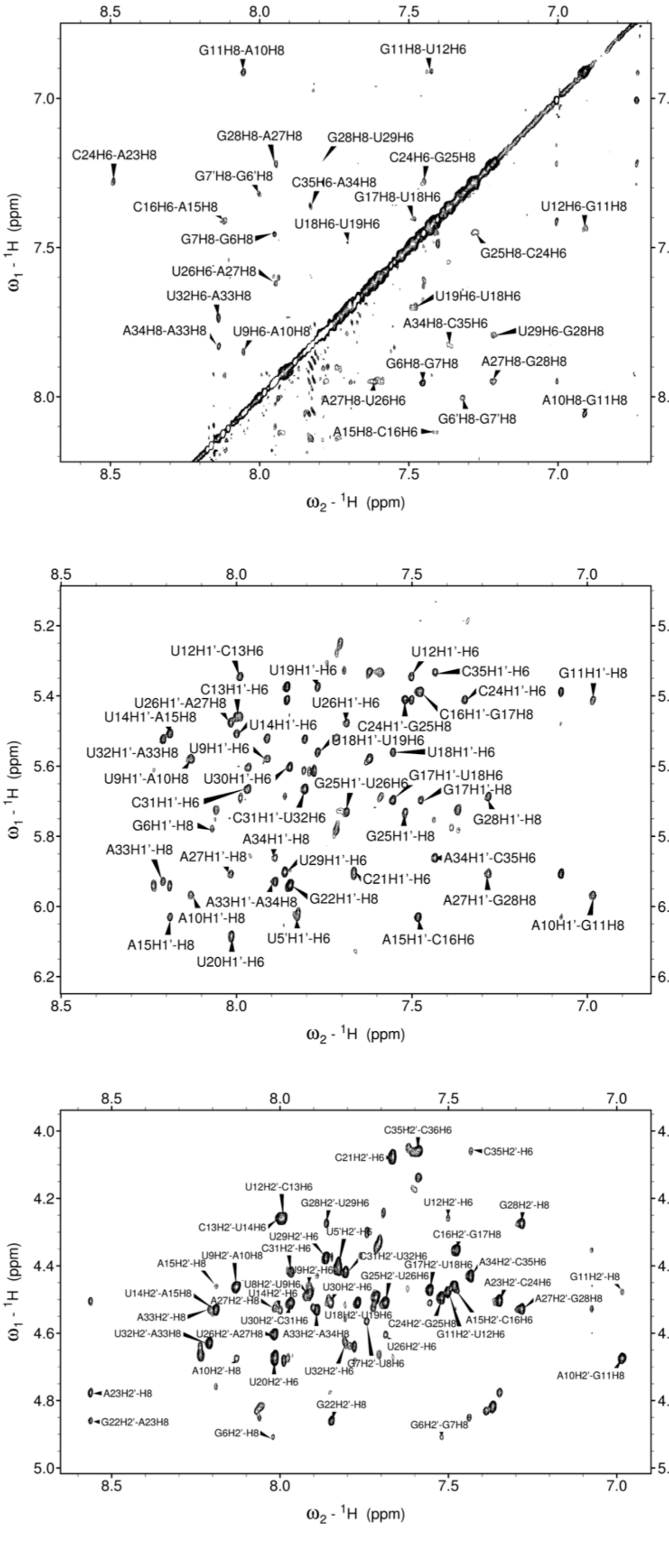


*Supplementary Figure S4.* NOESY spectra of non-exchangeable proton region (H6/H8-H6/H8 and sugar proton-H6/H8) for DenvBS. Deuteration of H5, H3’, H4’, H5’, H5’’ protons relieved the extensive overlap in the sugar proton region.

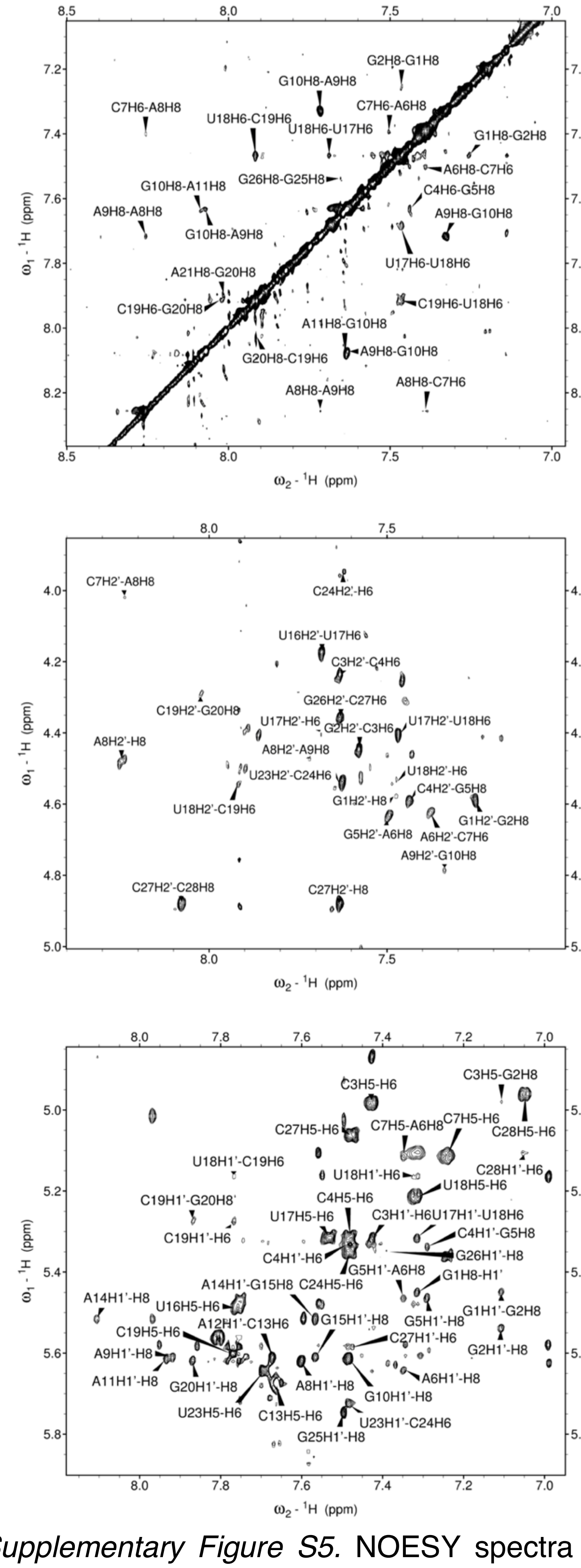


*Supplementary Figure S5.* NOESY spectra of non-exchangeable proton region (H6/H8-H6/H8 and sugar proton-H6/H8) for DenvTSL. Deuteration of H5, H3', H4', H5', H5'' protons relieved the extensive overlap in the sugar proton region.

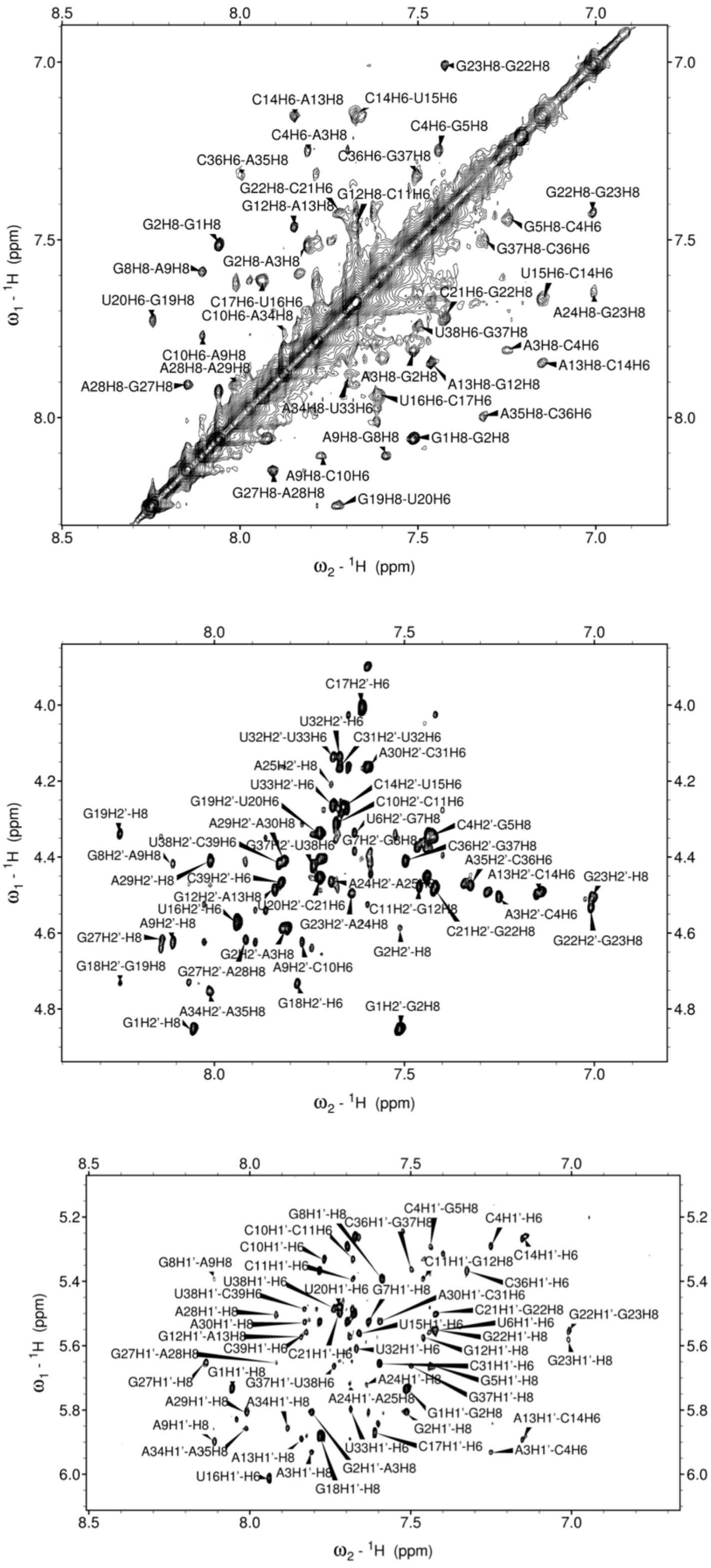


*Supplementary Figure S6.* NOESY spectra of non-exchangeable proton region (H6/H8-H6/H8 and sugar proton-H6/H8) for DenvSLAsh. Deuteration of H5, H3', H4', H5', H5'' protons relieved the extensive overlap in the sugar proton region.

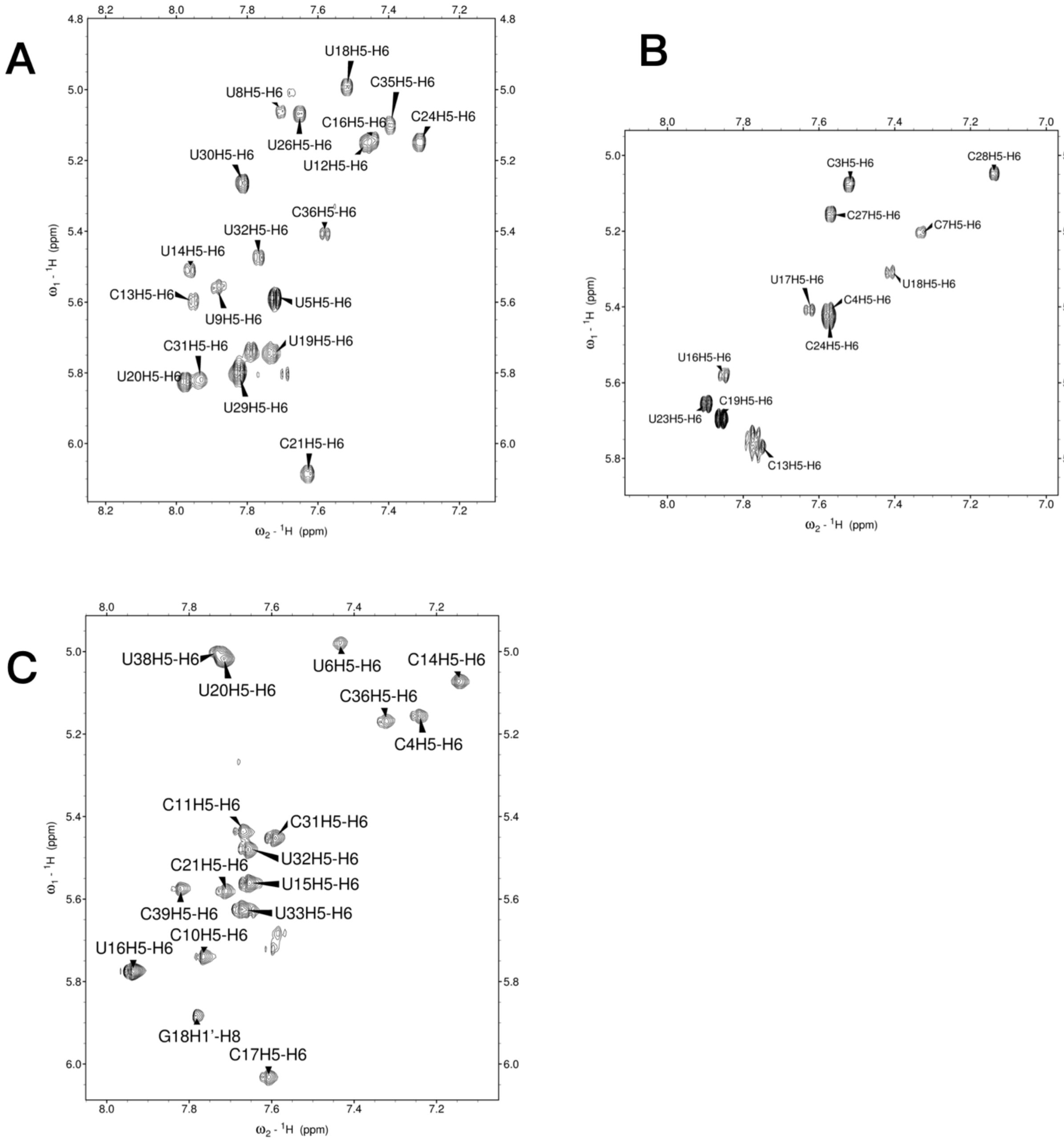


*Supplementary Figure S7.* TOCSY spectra for the three independently folded structural segments: (A) DenvBS, (B) DenvTSL, and (C) DenvSLAsh. The existence of a single monomeric conformation for each RNA was confirmed from the number of Ura and Cyt H5-H6 peaks in TOCSY spectra, which was as predicted for each sequence.

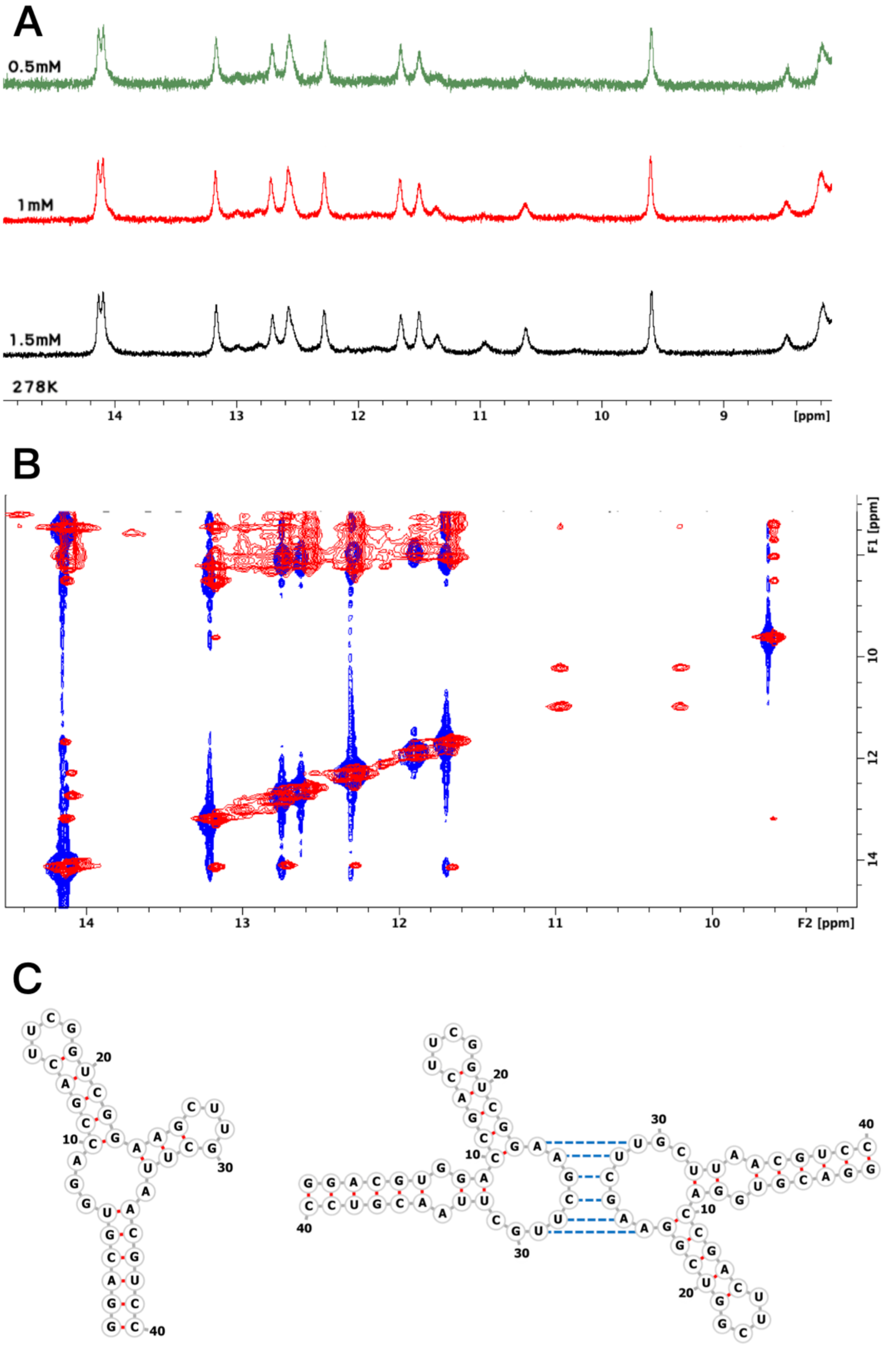


*Supplementary Figure S8.* (A) 1D ¹H NMR spectra of SLAshCUUG recorded at 5°C at concentrations of 0.5 mM (green), 1 mM (red), and 1.5 mM (black). Concentration-dependent peaks are observed between 10-12 ppm and between 12 and 13 ppm, which imply formation of new base pairs as a result of dimerization, which of course becomes more favorable at higher RNA concentration. (B) Overlay of 2D ¹H NOESY spectra for SLAshCUUG (red) and DenvSLAsh (blue). For SLAshCUUG, extra base pair peaks are observed between 10-11 ppm, consistent with formation of a GU base pair. By substituting CUUG with a GAAA tetraloop, the extra resonances are eliminated. (C) Secondary structures of the SLAshCUUG monomer (left) and the presumed SLAshCUUG dimer (right).

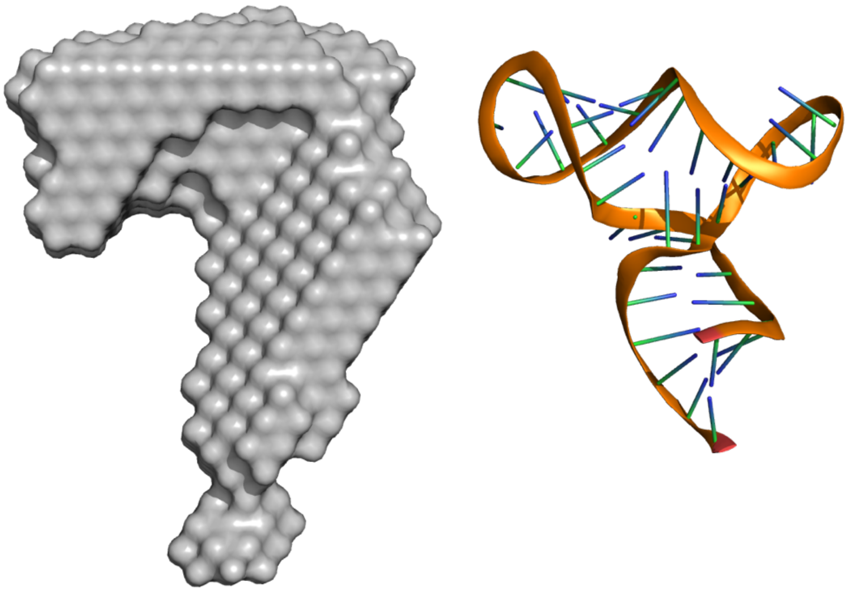

*Supplementary Figure S9*. The SAXS model of SLAshCUUG demonstrates a monomeric structure at concentrations below those used for NMR. Data was collected using the same process described in Method at RNA concentrations of 0.1-2 mg/mL. The structure of DenvSLAsh is shown on the right for comparison.

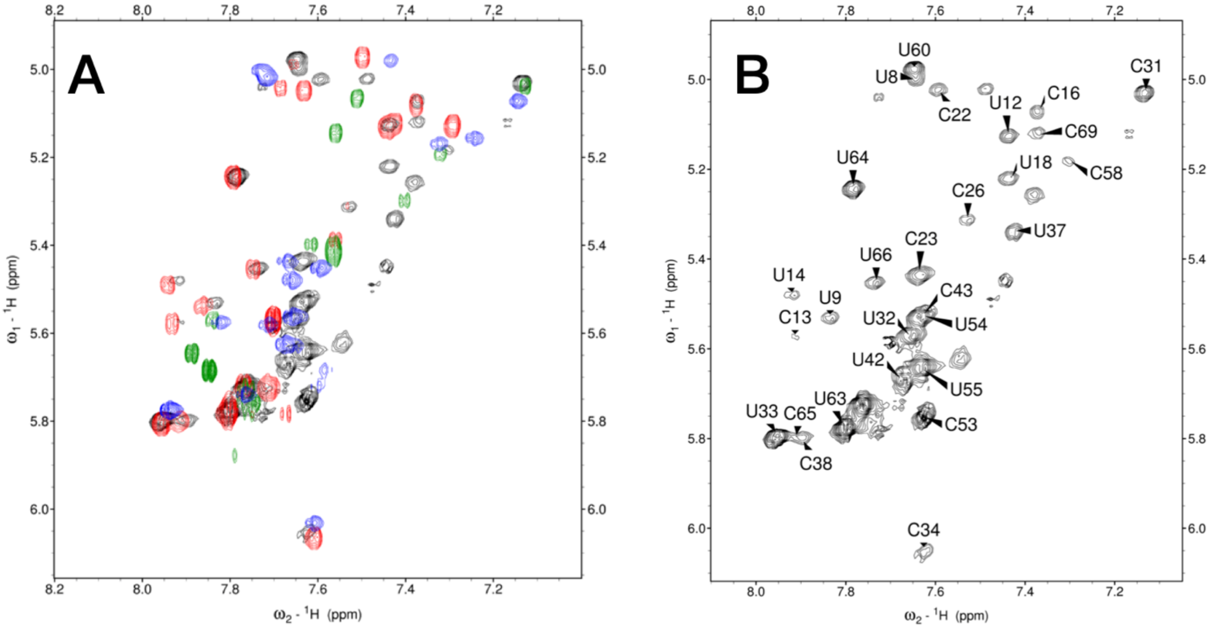


*Supplementary Figure S10.* (A) Overlay of the TOCSY spectra for DenvSLATL (black), DenvBS (red), DenvTSL (green) and DenvSLAsh (blue). The similarity of chemical shifts allowed us to assign H5-H6 cross-peaks for DenvSLATL, as shown. (B) H5-H6 peak assignments for the TOCSY spectrum of the complete DenvSLATL. A single dominant monomeric conformation was confirmed from the number of Ura and Cyt H5-H6 peaks in TOCSY spectra, which was consistent with its sequence.

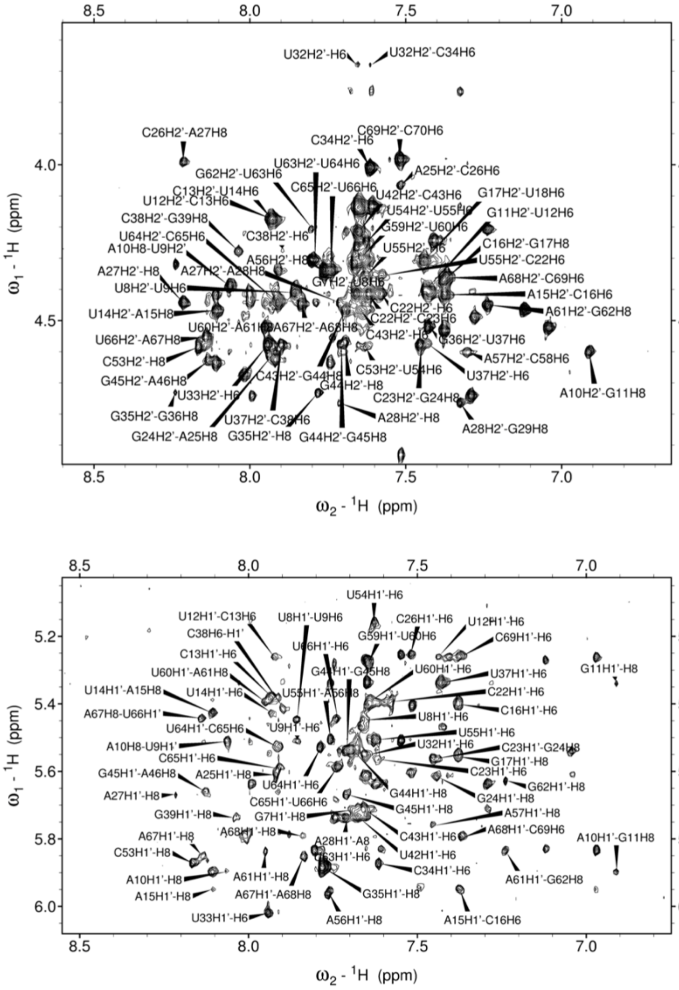


*Supplementary Figure S11.* NOESY spectra of the H1'-H6/H8 and H2'-H6/H8 region for DenvSLATL. Selective deuteration of H5, H3', H4', H5', H5'' protons was applied to relieve the extensive overlap in the sugar proton region, leading to interpretable and fully assignable spectra.

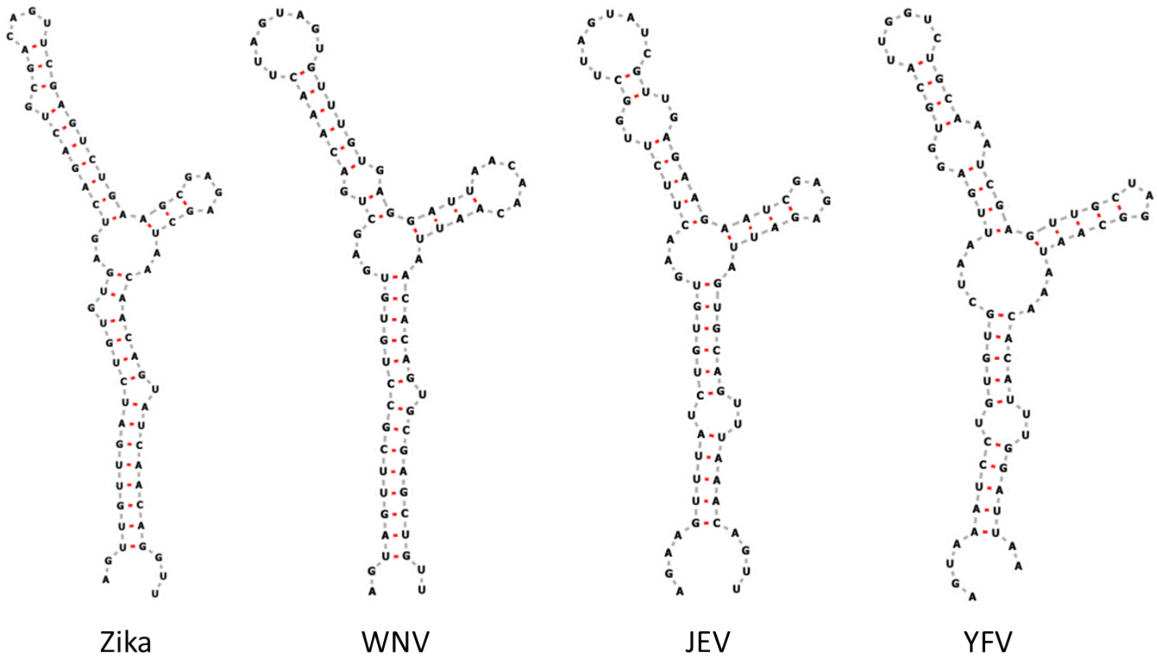


*Supplementary Figure S12.* Predicted SLA secondary structures, according to Vienna fold, for different flaviviruses (Lodeiro et al., 2009; Markham & Zuker, 2008; Zuker, 2003). Results are shown for Zika virus (NC_012532, nucleotide 1-106), West Nile virus (NC_001563, nucleotide 1-96), Japanese encephalitis virus (NC_001437, nucleotide 1-95), and Yellow fever virus (NC_002031, nucleotide 1-118).
